# Structural basis of SBDS dysfunction in Shwachman-Diamond Syndrome

**DOI:** 10.64898/2026.11.11.766026

**Authors:** Rajnandani Kashyap, Sinthyia Ahmed, Poonam Roshan, Edwin Antony, Sofia Origanti

## Abstract

Surveillance of functional centers in the mature 60S ribosomal subunit by SBDS licenses its translational debut. Timely release of eIF6 from 60S by SBDS is essential to permit 40S joining. Biallelic mutations in *SBDS* associated with Shwachman-Diamond syndrome (SDS), an inherited ribosomopathy, impair dissociation of eIF6 from 60S. How SBDS variants that bind to 60S structurally perturb the maturation and surveillance of 60S is yet to be deciphered. Here, we report high-resolution cryogenic electron microscopy (cryo-EM) structures of human SBDS and pathogenic variants in complex with human 60S. The structures reveal that disease mutations disrupt key interactions of SBDS with the functional centers of the 60S subunit destabilizing critical bases and protein interactions across the P-site, peptidyl transferase center, polypeptide exit tunnel and GTPase-associated center. These mutations remodel the N-terminal contacts with 60S, altering critical interactions with uL16 (RPL10) distant from the mutation site. Mutations in domains II and III trigger conformational changes that propagate across the entirety of SBDS and remodel distant ribosomal regions, revealing an elegant mechanism of pathogenic allostery. The profound effects of SBDS mutations on both the surveillance and mature conformation of the 60S subunit provide a structural framework for understanding the molecular pathology of SDS.

## Introduction

Eukaryotic ribosome biogenesis is a complex process involving more than 200 trans-acting assembly factors^1–6^. Cytoplasmic maturation of the 60S ribosomal subunit culminates with the release of eIF6^7–10^. eIF6 is an essential 60S assembly factor that also serves as a steric block to prevent the premature association of the 60S and 40S subunits^8,10–15^. Defects in maturation of 60S are marked by stalling of pre-60S subunits bound to eIF6 in the cytoplasm^8,9,12,16–18^. Rigorous surveillance mechanisms ensure that only accurately assembled 60S are fit for translation^8,9,12,16–18^. One such surveillance mechanism involves the Shwachman-Bodian-Diamond Syndrome protein (SBDS) that probes for the precise insertion of ribosomal protein 10 (RPL10 or uL16) during the late stages of 60S maturation^8,9,12,16,19^. Insertion of uL16 and appropriate positioning of its P-site loop supports completion of the peptidyl transferase center^18,20–24^. The final mature state of 60S is then probed by SBDS, which cooperates with EFL1-GTPase to release eIF6 and permit subunit interactions^8,9,12,19,25^.

eIF6 is bound to both pre-60S and mature free 60S subunits in the cell^7^. 60S subunits dissociated with peptidyl tRNA and nascent peptides as part of the ribosomal quality control (RQC) pathway are also bound by eIF6^26^, which likely prevents premature subunit joining. SBDS and EFL1-GTPase have been proposed to be minimal factors sufficient to release eIF6 from distinct states of 60S^7,8,12^. Given their significance, it is not surprising that factors associated with late-cytoplasmic 60S maturation are essential for growth and survival. Mutations in uL16 have been associated with ∼8% of pediatric T-cell acute lymphoblastic leukemias^17,27–30^. Disease mutations in uL16 inhibit release of eIF6 from the 60S subunit^17,18,21,22^. Similarly, eIF6 eviction is perturbed in Shwachman-Diamond Syndrome (SDS) that is predominantly associated with mutations in SBDS^8,9,12,16,19,31^. Loss of functional SBDS leads to subunit joining defects due to an inability to release eIF6^8,9,12,16,19^.

SDS is an inherited autosomal recessive disorder characterized by bone marrow failure, exocrine pancreatic insufficiency, and skeletal abnormalities with an increased risk for leukemias^12,32–39^. Majority of SDS patients carry biallelic mutations in SBDS^12,32^. The mutations are mostly compound heterozygous and associated with hypomorphic variants that lead to partial loss of function^12,32,40^. The common mutations are 183-184TA→CT, which introduces a stop codon at K62; splicing mutation 258+2T > C with 8bp deletion that leads to a truncated product by frameshift (C84fs3) and pathogenic missense mutations^12,32,38,41^.

Based on chemical shift perturbation analysis of in-solution NMR structures, SBDS mutations were broadly classified as either fold-destabilizing mutations, which impair SBDS stability and expression; or surface mutations which do not affect folding or stability but retain the ability to bind 60S^9,12^. X-ray and NMR-derived structures of archaeal and human SBDS indicate a three-domain architecture that is highly conserved^9,12,42–44^. The three domains of SBDS are: the N-terminal FYSH (Fungal, Yhr087wp, Shwachman) domain (Domain I; S2-S96), the central domain (Domain II; D97-A170), and the C-terminal domain (Domain III; H171-E250)^9,12^. A crosslinker-stabilized cryo-EM structure of human SBDS (4–5 Å) was solved by reconstitution of SBDS with pre-60S isolated from an SBDS mutant strain of *Dictyostelium discoideum*^8^. This structure provided key insights and showed domain I of SBDS nestled in the P-site and peptidyl transferase center (PTC) of the mature 60S^8^. Although side-chain interactions of SBDS with 60S could not be mapped due to resolution limitations, this cryo-EM structure, along with yeast studies on uL16 mutants and *in vitro* binding assays, were instrumental in deciphering the function of SBDS as a surveillance factor that proof-reads the mature 60S^8,9,12^.

How the 60S-surveillance activity of SBDS is affected by 60S-binding disease variants of SBDS is yet to be determined. Certain missense SBDS variants, including K33E and R126T, are expressed at levels comparable to wild type and exhibit 60S-binding activity^12,16,40^. Missense mutants predominantly cause subunit joining defects^12,16^. Many of these mutants are pathogenic in mice and other organisms^19,35,36,45–47^. Mice carrying *Sbds^R^*^126^*^T/R^*^126^*^T^* or *Sbds^R^*^126^*^T/-^* alleles do not survive birth, further highlighting the significance and impact of missense variants on SBDS function^36,46^. 60S-binding SBDS mutants have been shown to inhibit the GTPase activity of EFL1, likely by disrupting its recruitment^8,9,48,49^. However, the extent to which these mutants perturb EFL1 GTPase activity and 60S ribosomal subunit levels appears to vary, suggesting that individual 60S-binding SBDS mutants exert distinct functional effects^8,9,48–50^. SBDS defects in SDS are intrinsically associated with the disruption of eIF6 release. However, releasing eIF6 does not fully restore the hematopoietic defects, suggesting that SBDS mutants may have a larger role in impairing 60S maturation^12,16^.

The footprint of SBDS on 60S spans ∼100 Å, raising the fundamental question of how mutations positioned across the different domains in SBDS impair surveillance and subunit joining. To address this question, we determined high-resolution single-particle cryo-EM structures of human wildtype (WT) SBDS and pathogenic variants in complex with the human 60S ribosomal subunit. Three functionally characterized missense variants harboring mutations in each of the three domains of SBDS were interrogated. SBDS acts as a long-range allosteric scaffold that can simultaneously communicate across multiple functional centers of the mature 60S subunit spanning the P-site, PTC, polypeptide exit tunnel and GTPase-associated center. Mutations in domains II and III destabilize key protein and base interactions through long-range allosteric communication propagated across all domains of SBDS bound to 60S. These SBDS-60S complexes capture the first structural snapshots of disease mutants associated with the inherited ribosomopathy-Shwachman-Diamond Syndrome and highlight a distinct allosteric mechanism that contributes to SDS pathology.

## Results

### Cryo-EM structures of SBDS-WT and disease variants bound to human 60S

Given the crucial role of SBDS in surveillance of 60S maturation, we aimed to determine the effects of SBDS variants on the surveillance mechanism. SBDS-WT, and SBDS variants with single site mutations in each of the three domains were purified: K33E, R126T and R175W (**Supplementary Fig. 1A**). SBDS associates with 60S independent of EFL1. Consistent with previous observations with K33E and R126T^16^, the missense variants also bound to 60S similar to WT in an *in vitro* binding assay (**Supplementary Fig. 1B**). Circular dichroism (CD) profiles of SBDS proteins and mutants (not bound to 60S) show no quantifiable differences in the spectral shape suggesting no gross structural changes in solution (**Supplementary Figs. 1C-F**). However, there is a distinct reduction in the amplitude of CD signals for SBDS-K33E and SBDS-R175W suggesting a reduction in secondary structure content. Thermal denaturation CD profiles of SBDS proteins indicate a -5 to -10 °C *ΔT_m_* shift for the three mutants compared to SBDS-WT suggesting differences in conformational stability (**Supplementary Figs. 1G & H**). These data suggest that the mutant SBDS proteins can assess differential conformational states when not bound to 60S. From a functional standpoint, these changes could be pronounced in the context of 60S, but such studies have never been performed. To address this void, we reconstituted human SBDS (WT or mutants) with human 60S and solved their structures using single particle Cryo-EM (**Fig. 1**).

**Figure 1.**
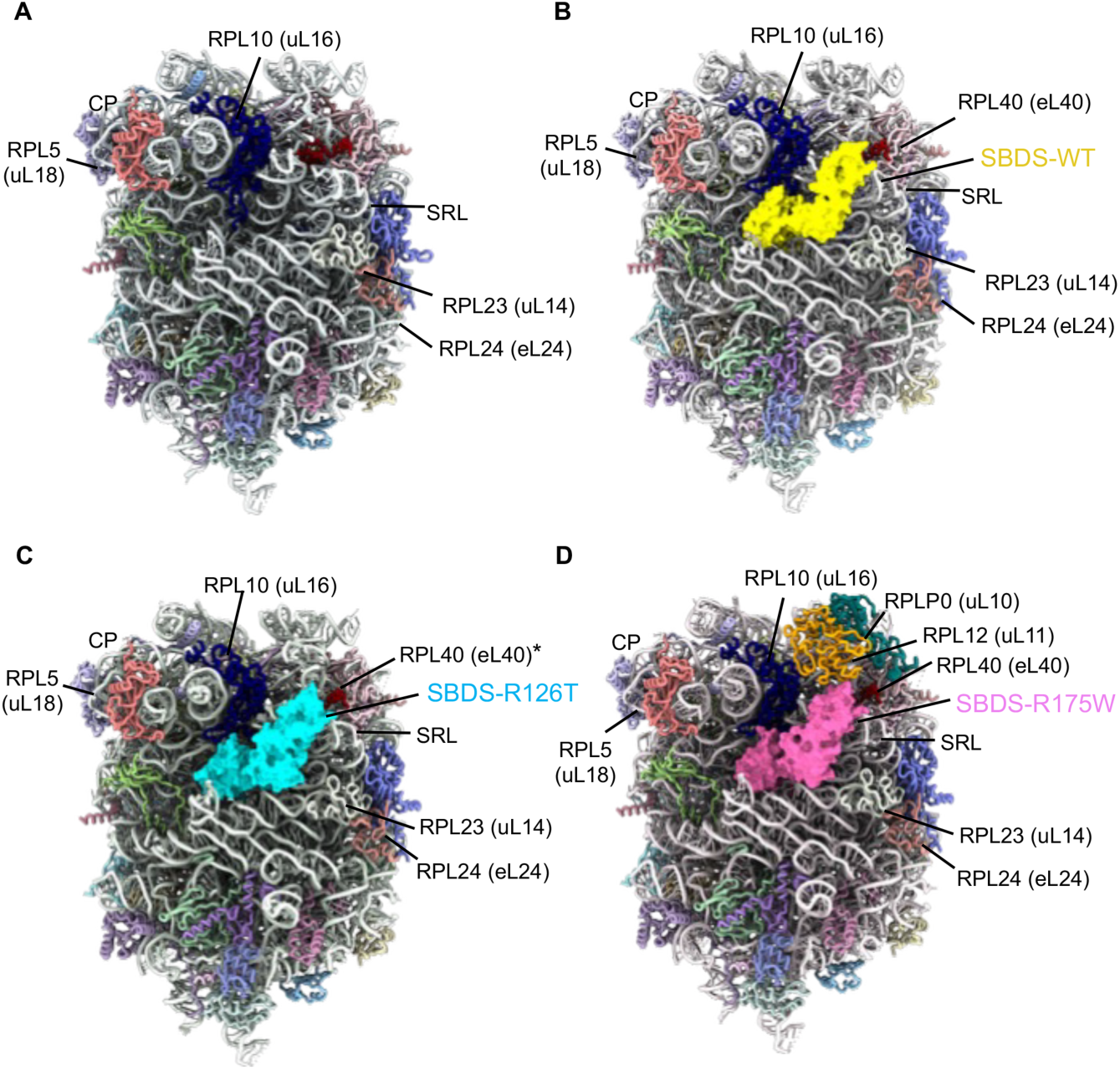
Cryo-EM structures of SBDS-bound human 60S ribosomal subunits. Shown are the structures of *(A)* the vacant human 60S subunit (2.49 Å), *(B)* the SBDS-WT-bound 60S complex (2.92 Å), *(C)* the SBDS-R126T-bound 60S complex (3.17 Å), and *(D)* the SBDS-R175W-bound 60S complex (3.09 Å). SBDS is displayed as a surface representation, whereas ribosomal proteins are shown as individually colored cartoon models. *eL40 is modeled in the R126T structure as it displays sparse density.

We solved the structures of vacant human 60S (2.49 Å; **Fig. 1A & Supplementary Fig. 2**), and in complex with SBDS-WT (2.92 Å; **Fig. 1B & Supplementary Fig. 3**), SBDS-R126T (3.17 Å; **Fig. 1C & Supplementary Fig. 4**), or SBDS-R175W (3.09 Å; **Fig. 1D & Supplementary Fig. 5**). Overall, the structures showcase the highly conserved three-domain architecture of SBDS for both WT and mutants. Although the three domains of SBDS are intrinsically flexible, neither the R126T or R175W mutation alters the global positioning of the three domains on 60S. Modeling eIF6 onto our structures indicates that SBDS adopts a more distal position relative to eIF6 and lacks direct SBDS-eIF6 contacts (**Supplementary Figs. 6A & B**), in agreement with previous report^8^.

Despite five separate attempts (25,400 micrographs), and scouring through 1.31 million 60S particles, we were unable to solve a structure of SBDS-K33E (domain I mutant) (**Supplementary Fig. 7**). From this large dataset we were able to extract 49,359 particles that possessed sparse density for SBDS-K33E on 60S (**Supplementary Figs. 7 & 8**). The empty 60S particles from these grids were of high resolution with excellent views and we could readily arrive at a solvable 3D volume (**Supplementary Fig. 8A**). In contrast, the SBDS-K33E bound 60S particles suffered from severe orientation bias (**Supplementary Fig. 8B**). The disparity in behavior of the empty 60S versus the SBDS-K33E-60S particles suggest overall structural changes and possible air-water-interface effects introduced by this domain I mutant of SBDS. Whether this is a common feature of other known SBDS domain I mutations remains to be investigated.

In our structures (**Figs. 2A-D & Supplementary Fig. 9**), SBDS occupies a large open cleft on 60S and contacts Helix-89 (H89), Helix-93 (H93) and the conserved sarcin-ricin loop (SRL) of rRNA. Domain III of SBDS is proximal to the sarcin-ricin loop of rRNA, which is important for activation of translational GTPases. Domain III of SBDS is also proximal to eL40 (RPL40) that supports the stable incorporation of uL16 (RPL10) and lies in the GTPase-associated center. Overall, there are three major structural differences between the complexes: SBDS-R175W is the most ordered of the three and results in subsequent stabilization of both uL11 (RPL12) and uL10 (RPLP0; **Fig. 2D**) which are absent in the vacant 60S, SBDS-WT and SBDS-R126T bound 60S structures (**Figs. 2A-C**). uL11 interacts with uL10 (RPLP0) that forms the base of P-stalk. The dynamic P-stalk is completed by assembly of the P1 and P2 heterodimers bound to uL10. However, we did not observe any density for the P1/P2 heterodimers in any of our structures. In addition, the L1 stalk could not be resolved due to poor density. SBDS-R126T-60S complex is the least ordered and does not show strong density for eL40 unlike vacant 60S or SBDS-WT or SBDS-R175W-60S structures (**Supplementary Fig. 10**). Finally, the N-terminus of human SBDS occupies the P-site with the first 15 residues interacting with key residues in uL16 (**Fig. 2E & Supplementary Fig. 11**), an integral part of the PTC that aids in positioning tRNAs in the PTC. While this region makes key contacts with several rRNA bases (detailed below) in both the SBDS-WT and SBDS-R175W-bound 60S complexes, it is disordered with very sparse density in the SBDS-R126T-60S structure (**Fig. 2E & Supplementary Fig. 11**).

**Figure 2.**
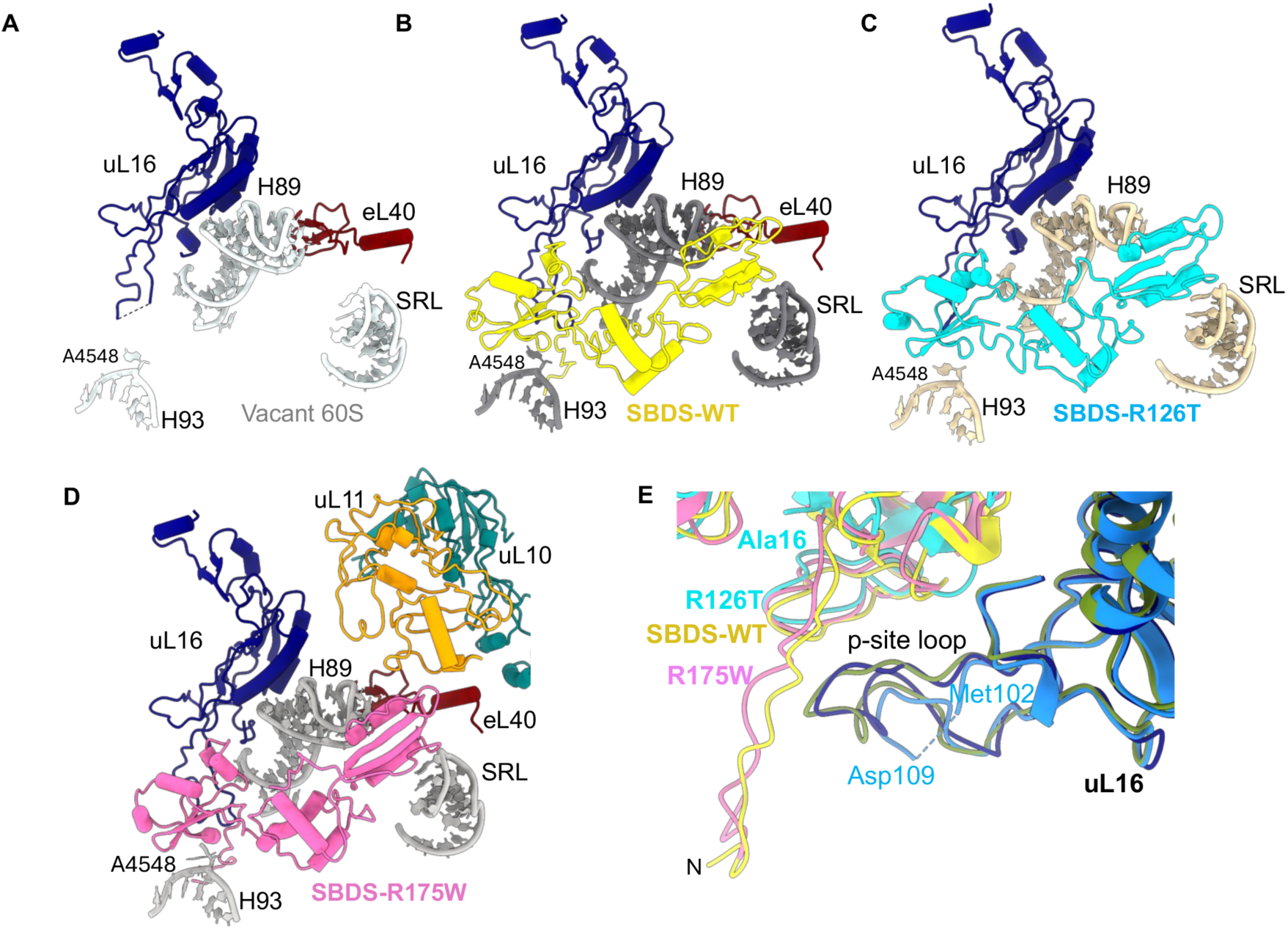
Positioning of SBDS on the ribosomal P-site. *(A)* Location of helix 89 (H89), helix 93 (H93), and the sarcin-ricin loop (SRL) on the vacant human 60S subunit. Structures of *(B)* SBDS-WT, *(C)* SBDS-R126T, and *(D)* SBDS-R175W bound to the 60S subunit. SBDS and uL16 (blue) positions on the 60S subunit are similar in all three complexes. Ribosomal protein eL40 (reddish brown) is resolved in the SBDS-WT- and SBDS-R175W- bound 60S structures but is poorly resolved in the SBDS-R126T-bound 60S complex. Ribosomal proteins uL11 (orange) and uL10 (teal) are observed only in the SBDS-R175W-bound 60S structure. The position of nucleotide A4548, which forms a key contact with domain I of SBDS, is indicated. *(E)* Superposition of SBDS-WT, SBDS- R126T, and SBDS-R175W highlighting differences in the N-terminal region of SBDS. The N-terminal segment is resolved in the SBDS-WT- and SBDS-R175W-bound 60S structures but is not observed in the SBDS-R126T- bound complex.

### SBDS mutations destabilize the P-site loop of uL16 and disrupt PTC surveillance

Stable incorporation of uL16 and insertion of its essential P-site loop (aa102 to 112) is crucial to complete the PTC^23^. During maturation, uL16 incorporation is facilitated by the nuclear export factor NMD3. Conformational rearrangements in H89 and H38 mediated by NMD3 create a cleft for the stable binding of uL16, which in turn releases NMD3^23^. Proper positioning of the uL16 P-site loop within this cleft is critical, as deletion of the loop or mutations within the loop, inhibit 60S subunit maturation and disrupt subunit joining^17,18^. In our vacant 60S structure, the uL16 loop is dynamic and displays sparse density (**Fig. 3; Supplementary Fig. 12A**). However, upon binding to SBDS-WT, the uL16 loop is stabilized and shows key interactions with the N-terminal residues of SBDS (**Fig. 2E and 3B; Supplementary Fig. 11 and 12B**). In contrast, although we have good resolution for most of SBDS-R126T, density for the N-terminal 15 residues is sparse suggesting that it is more dynamic in this mutant (**Fig. 2E and 3C; Supplementary Fig. 11 and 12C**). Correspondingly, sparse density was observed for the uL16 loop (aa 102 to 109) for the SBDS-R126T mutant (**Supplementary Fig. 11B**). This was not an artifact due to lack of protein stability as mass spectrometric and western blot analysis confirmed that the N- and C-terminal residues of SBDS-R126T are intact (**Supplementary Fig. 13**). These data further suggest that the stable anchoring of the N-terminal residues of SBDS reciprocally stabilizes the P-site loop of uL16.

**Figure 3.**
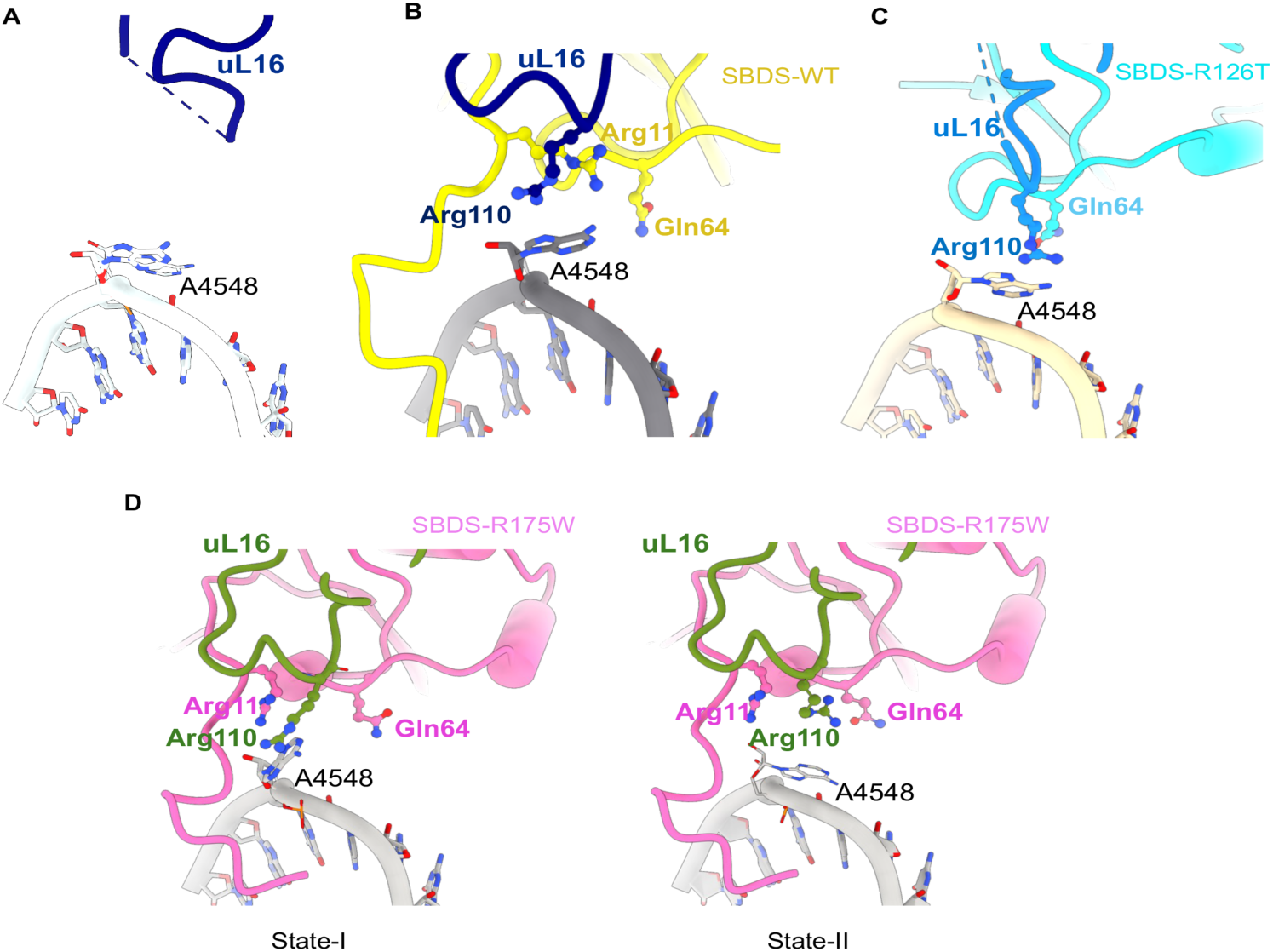
SBDS stabilizes interactions of Arg110 of uL16 with A4548 base of the peptidyl transferase center. *(A)* Structure of the vacant human 60S subunit showing the conformation of A4548 base within the peptidyl transferase center. Close-up views of the A4548 base interface in the *(B)* SBDS-WT, *(C)* SBDS-R126T, and *(D)* SBDS-R175W-bound 60S structures. In the SBDS-WT-60S complex, Arg11 is positioned between Gln64 and Arg110 (uL16), acting as a spacer within the SBDS-H93 interaction network, which stabilizes interactions of Arg110 with A4548. In the SBDS-R126T and SBDS-R175W-60S complexes, Arg11 is displaced from this position, allowing Arg110 to interact with Gln64. These rearrangements are accompanied by changes in the conformation of nucleotide A4548. Two distinct conformational states of A4548 are captured in the cryo-EM reconstructions for the R175W-bound 60S complex. The alternative A4548 conformations correlate with remodeling of the Arg11-Gln64-Arg110 interaction network at this interface.

Similar to SBDS-WT, the SBDS-R175W mutant also shows stable interactions with the uL16 loop (**Fig. 2E and 3D**). However, key interactions of the uL16 loop with the rRNA are perturbed in this mutant. In yeast, stable insertion of the uL16 loop promotes side chain interactions of a key conserved Arg110 residue with the A2971 base of 25S rRNA that marks the completion of the peptidyl transferase center^23^. A2971 in yeast (A2602 in *E. coli* and A4548 in human) is required for proper positioning of the A-site and P-site tRNA 3′CCA ends and plays essential roles in peptide release^23,51^. SBDS-WT stabilizes the flipped-out conformation of A4548 through π-stacking interactions with Arg110 in the P-site loop of uL16 and assesses the integrity of this crucial base (**Fig. 3B**). Arg11 is adjacent to Gln64 in SBDS-WT and positioned as a spacer to orient Arg110 (uL16) away from Gln64 (SBDS-WT). This spacing allows Arg110 of uL16 to interact with A4548. In contrast, in the SBDS-R126T mutant, the absence of the N-terminal region carrying Arg11 results in a loss of proper spacing leading to conformational rearrangement and direct H-bonding between Arg110 and Gln64. This results in loss of A4548 sensing by Arg110 (**Fig. 3C**). The SBDS-R175W mutant displays two different conformations for A4548 and Arg110 (**Figs. 3D; Supplementary Figs. 12D & E**). In both states, Arg11 in SBDS fails to slot in as a spacer leading to Arg110 in uL16 sampling two rotational states (∼60°) and failing to stabilize A4548. This results in a ∼90° flipping of A4548 between the two states which in turn functionally results in disruption of PTC proofreading activity. These results suggest that both SBDS-R126T and SBDS-R175W mutants markedly disrupt the interrogation of the PTC by altering the π-stacking interactions of the A4548 base with the P-site loop of uL16 in the PTC.

Ser104, located within the uL16 loop, is important for ribosomal transitions from a non-rotated to rotated state^20^. Ribosomes are usually in the classical non-rotated state, but they transition to a rotated/hybrid state during the elongation cycle where the tRNAs are positioned in A/P and P/E hybrid states^20^. A Ser104 to Asp substitution shifts the ribosome to a rotated state and decreases mRNA decoding accuracy of the ribosome^20^. Intriguingly, SBDS has been proposed to favorably bind to the non-rotated state and stabilize this state through interactions with the P-site^20^. How SBDS contributes to the maintenance of this classical state remains unclear. We are missing density for the loop containing Ser104 in uL16 in vacant 60S structure (**Fig. 4A; Supplementary Fig. 14A)**. Intriguingly, SBDS-WT does not display any side chain interactions with Ser104 in the uL16 loop (**Fig. 4B; Supplementary Fig. 14B**). The lack of density for this region in SBDS-R126T complex precluded assessment of Ser104 interactions (**Fig. 4C; Supplementary Fig. 14C**). In contrast, the SBDS-R175W mutant establishes a continuous interaction network that links domain I to the uL16 loop. At the center of this network, Gln52 hydrogen bonds with both Thr13 and Ser104, creating a Thr13-Gln52-Ser104 hydrogen bonding triad that stabilizes the orientation of Gln52 and promotes persistent engagement with the uL16 loop (**Fig. 4D; Supplementary Fig. 14D**). This hydrogen-bonding network is further reinforced by a second set of interactions involving Lys33, Asn14, and Asp97. Specifically, Lys33 forms electrostatic and polar contacts with Asp97 and Asn14, generating a stabilizing interaction network that rigidifies domain I. Together, these coupled hydrogen-bonding and electrostatic interaction networks lock the SBDS-R175W mutant into a more stable conformation, positioning Gln52 for sustained interaction with Ser104 and thereby reducing the conformational flexibility of the uL16 loop. The absence of these stabilizing interaction networks in SBDS-WT and SBDS-R126T suggests that the enhanced rigidity observed for R175W is unique to the mutant. Two nucleotides - U4531 (U2585 in *E. coli* 23S rRNA) and U4452 (U2506 in *E. coli* 23S rRNA), are also integral to the PTC and must be accurately positioned for binding of tRNA and peptide release^51–56^. They are often targeted by arrest peptides to render ribosomes inactive^52,53,57^. In the vacant 60S structure, these nucleotides face each other and are in a frayed conformation relative to WT-SBDS-bound 60S (**Fig. 5A; Supplementary Fig. 15A and B**). U4531 and U4452 positioning in the SBDS-R126T mutant structure looks similar to the vacant 60S (**Figs. 5A & B; Supplementary Fig. 15C**). Insertion of the N-terminal six residues of SBDS-WT and SBDS-R175W crowds this pocket and restrains the conformation of both nucleotides (**Figs. 5 A, B & C; Supplementary Fig. 15B & D**). These findings suggest that the dynamic nature of the N-terminus of SBDS-R126T is unable to modulate the conformational positioning of U4531 and U4452.

**Figure 4.**
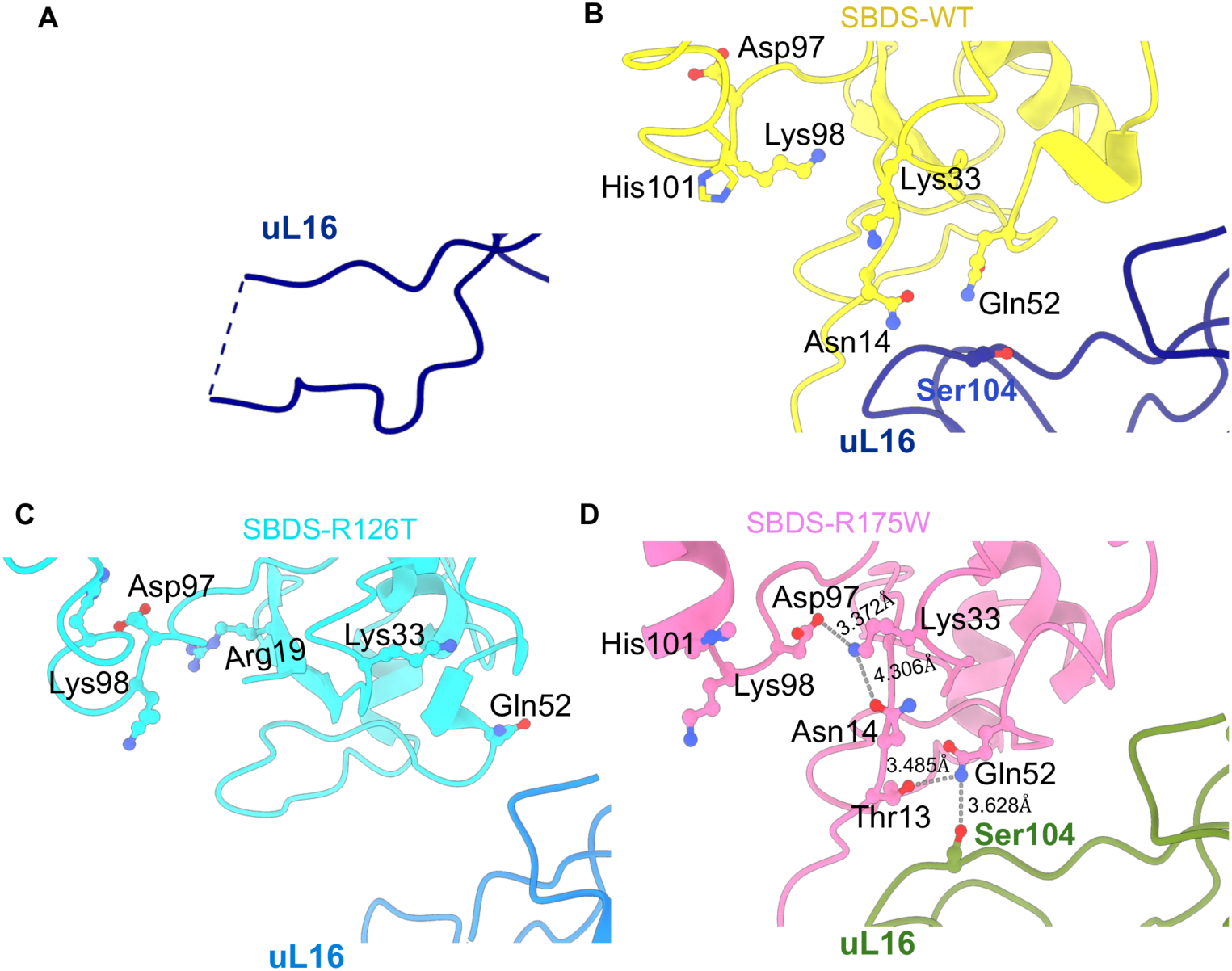
SBDS-R175W stabilizes the uL16 loop and interacts with the key Ser104 residue through coupled intramolecular interaction networks. *(A)* Ser104 within the uL16 loop is disordered in the vacant 60S structure. *(B)* In the SBDS-WT complex, Ser104 is resolved but does not form direct side-chain interactions with SBDS. *(C)* In the SBDS-R126T complex, disorder of the uL16 loop precludes assessment of interactions involving Ser104. *(D)* In the SBDS-R175W complex, Gln52 forms hydrogen-bonding interactions with Thr13 and Ser104, generating a Thr13-Gln52-Ser104 triad that links SBDS domain I to the uL16 loop. Additional interactions involving Lys33, Asn14, and Asp97 further stabilize domain I.

**Figure 5.**
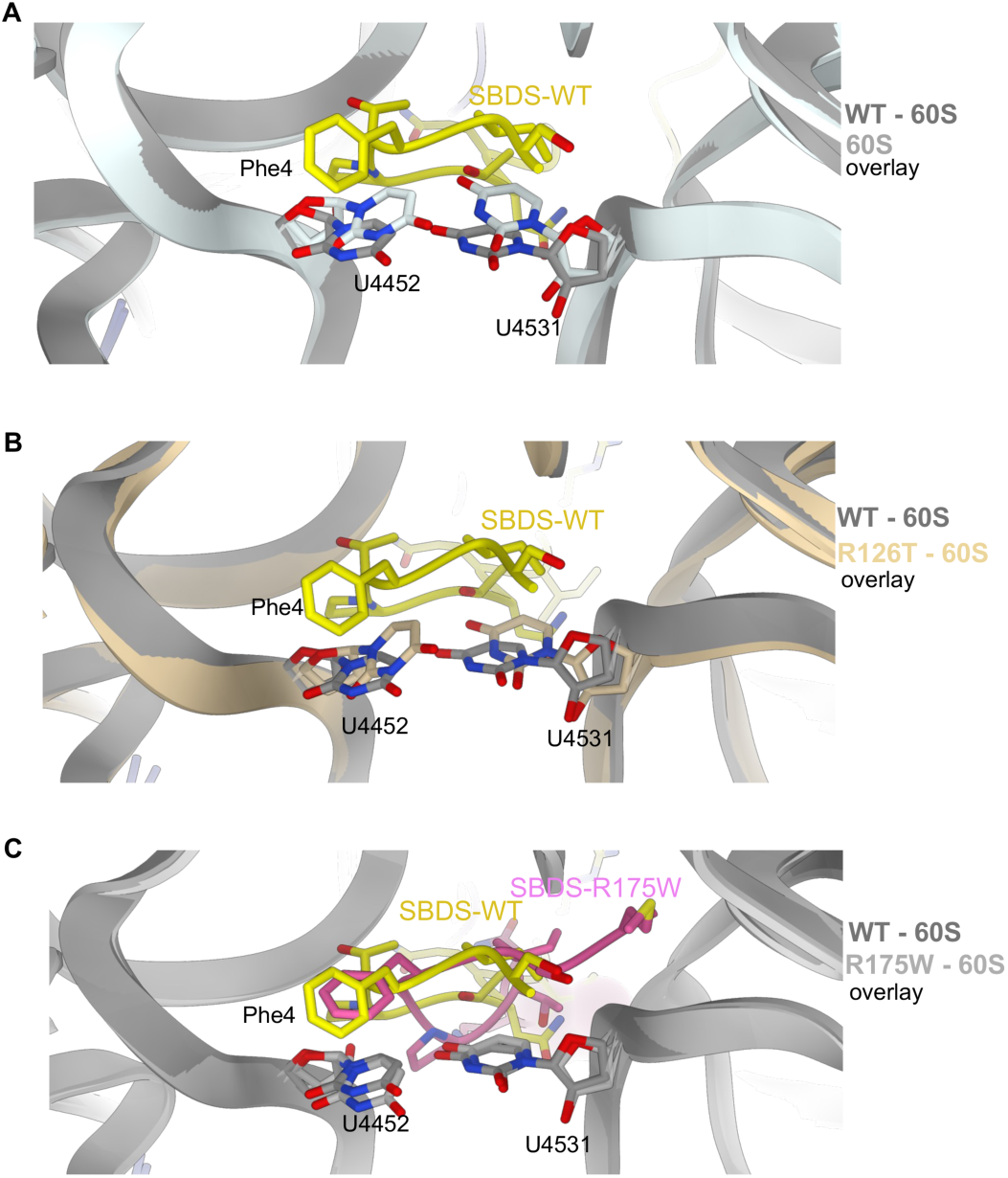
Differential effects of SBDS disease mutants on the positioning of nucleotides U4452 and U4531 in the peptidyl transferase center. *(A)* The N-terminus of SBDS occupies a pocket adjacent to U4452 and U4531, two highly conserved PTC nucleotides, shifting their conformations relative to the vacant 60S state and restricting their positional variability. *(B)* In the R126T mutant, the N-terminus fails to occupy this pocket, resulting in nucleotide conformations that more closely resemble the vacant 60S state and increased conformational flexibility of both PTC residues. *(C)* Although repositioned relative to WT, the N-terminus of SBDS-R175W remains within the pocket surrounding U4452 and U4531, maintaining a local nucleotide arrangement similar to that observed in the SBDS-WT-60S complex.

### SBDS variants perturb surveillance of the polypeptide exit tunnel

The N-terminal residues of SBDS are inserted into entrance of the polypeptide exit tunnel (PET). The PET is a structural conduit through which nascent polypeptides pass through as they exit from the 60S. A3908 (A2062 in *E. coli*) is a flexible base that gates the tunnel such that its positioning into the tunnel or towards the tunnel wall can either constrict the tunnel or provide space for peptide exit^51,58^. In the vacant 60S structure, A3908 is positioned away from the tunnel lumen (**Fig. 6A; Supplementary Fig. 16A**) and binding of SBDS-WT does not alter the conformation of this base (**Fig. 6B; Supplementary Fig. 16B**). However, A3908 is rotated by ∼80° into an alternate conformation such that it projects into the lumen in the presence of the SBDS mutants (**Figs. 6C-E; Supplementary Figs. 16C & D**). For the SBDS-R175W mutant, Phe4 stabilizes the rotated state of A3908 through π-stacking interactions (**Fig. 6D & E**). While we are unable to assess Phe4 interactions for the SBDS-R126T mutant due to the dynamic nature of the N-terminus, it is possible that similar interactions could orient A3908 in this intermediate flipped state. These results suggest that the mutants disrupt the conformational dynamics of A3908 and stabilize an intermediate state that could constrict the exit tunnel and likely affect proper positioning of the C-terminus of Reh1, a protein associated with the late-stage 60S maturation in the tunnel lumen in yeast^59^.

**Figure 6.**
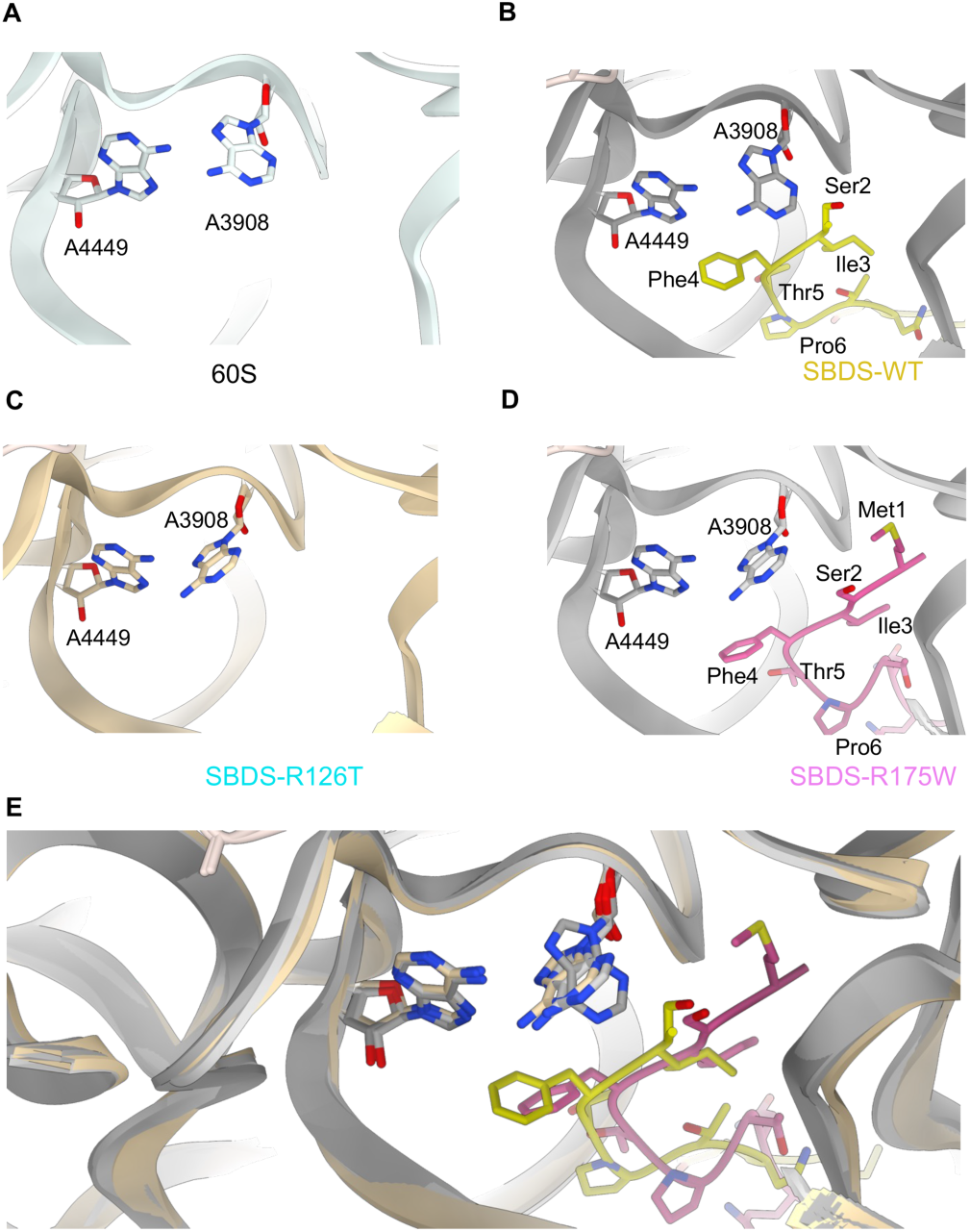
Conformation of the polypeptide exit tunnel gate nucleotide A3908 is modulated by mutations in SBDS. *(A)* A3908 adopts a conformation that is positioned closer to the polypeptide exit tunnel wall in the vacant 60S subunit. *(B)* Binding of SBDS-WT does not alter the orientation of A3908, *(C)* In the SBDS-R126T complex, A3908 is rotated into an alternative conformation relative to the vacant 60S and SBDS-WT structures and projects into the tunnel lumen. *(D)* In the SBDS-R175W complex, A3908 adopts a similar rotated conformation that is potentially stabilized by π-stacking interactions with Phe4. *(E)* Superposition of the vacant 60S, SBDS-WT, SBDS-R126T, and SBDS-R175W structures highlighting the repositioning of A3908 in the mutant complexes. A4449 is depicted as a reference for a stable (immobile) nucleotide.

### Effects of SBDS variants on the GTPase-associated center

Domain III of SBDS binds to the GTPase-associated center (GAC) of the 60S subunit. Both the P-stalk and SRL are integral parts of the GAC and aid in the docking and activation of translational GTPases^60–67^. A4605 is a key base in the SRL loop that is targeted by toxins such as ricin leading to an inhibition of eEF2 during elongation (**Fig. 7A; Supplementary Fig. 17A**)^66,68^.

**Figure 7.**
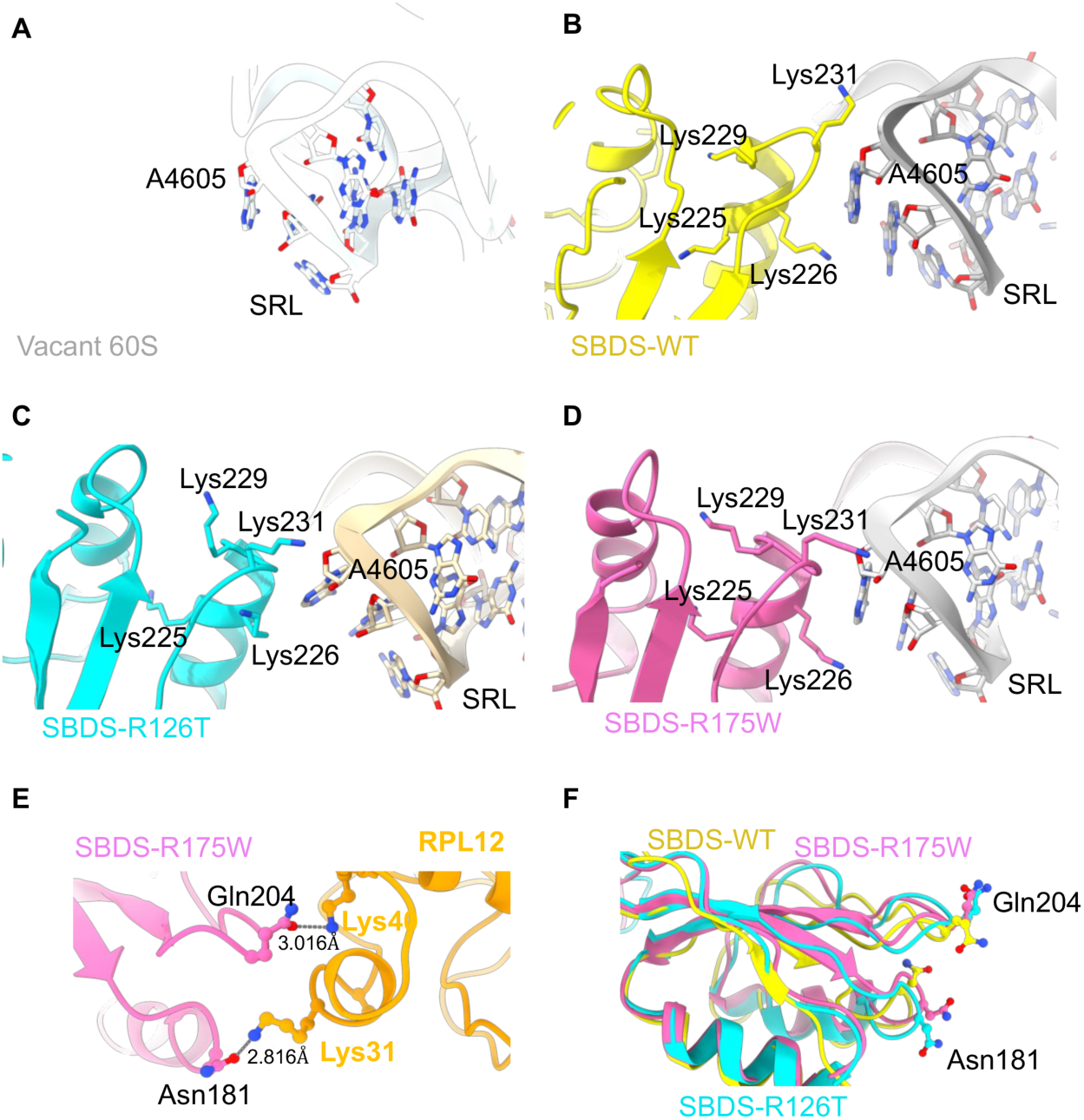
SBDS mutations remodel contacts between domain III and key regions of the GTPase-associated center. *(A)* Position of nucleotide A4605 in the SRL of the vacant 60S structure. *(B)* In the SBDS-WT structure, Lys231 of domain III is oriented away from the key conserved sarcin-ricin loop (SRL) nucleotide A4605. In the *(C)* SBDS-R126T and (*D)* SBDS-R175W structures Lys231 is repositioned proximal to A4605. *(E)* In the SBDS-R175W structure, Asn181 and Gln204 of domain III interact with Lys31 and Lys40 of RPL12, respectively, at the P-stalk base. *(F)* Superposition of the SBDS-WT, SBDS-R126T, and SBDS-R175W structures highlighting the relative positions of Asn181 and Gln204. Gln204 adopts a distinct orientation in the SBDS-R175W structure relative to SBDS-WT and SBDS-R126T that is compatible for interaction with RPL12.

**Figure 8.**
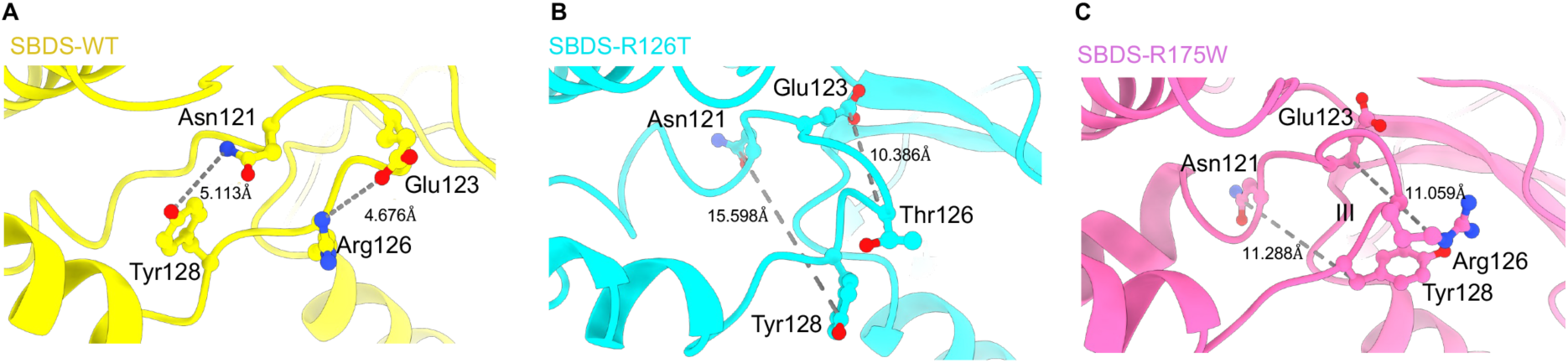
Structural communication across SBDS domains propagates the effects of disease mutations. *(A)* In the SBDS-WT structure, Arg126 forms side-chain interactions with Glu123 and positions Asn121 in close proximity to Tyr128, establishing a local interaction network within domain II. *(B)* In the SBDS-R126T structure, substitution of Arg126 disrupts interactions with Glu123 and increases the separation between residues that form the domain II interaction network in the WT structure. *(C)* In the SBDS-R175W structure, despite the mutation being located in domain III, the interaction network involving Glu123, Asn121, and Tyr128 is similarly disrupted, resulting in longer inter-residue distances relative to SBDS-WT.

Lys231 in domain III of SBDS-WT is positioned away from A4605 (**Fig. 7B; Supplementary Fig. 17B**). However, for both the mutants, the side chain of Lys231 is proximal to A4605 (**Fig. 7C & D; Supplementary Fig. 17C & D)**. R126T and other SBDS mutants are known to inhibit the GTPase activity of EFL1^8,9^. This suggests that the side chain interactions of Lys231 of mutants with SRL could prevent the docking and activation of GTPases by the mutants. In addition, only SBDS-R175W shows electrostatic interactions with RPL12 which is anchored along with RPLP0 to form the P-stalk base. Asn181 and Gln204 of SBDS-R17W interact with Lys31 and Lys40 of RPL12, respectively (**Fig. 7E; Supplementary Fig. 17E**). However, compared to both SBDS mutants, Gln204 is differentially positioned in the SBDS-WT structure suggesting that these residues are likely unable to interact with RPL12 and therefore, are unable to stably anchor the dynamic P-stalk (**Fig. 7F**). This suggests that the SBDS-WT does not interfere with the conformational flexibility of the P-stalk needed for GAC. However, the R175W mutant severely restricts conformational dynamics of the P-stalk. This structural property likely explains the functional defect in the SBDS-R175W mutant.

### Allosteric communications across domains propagate mutational effects

SBDS occupies a rather large footprint on 60S that spans ∼100 Å (**Fig. 1**). The N-terminus of SBDS that engages the PTC is situated over ∼60-80 Å from domains II and III where the R126T and R175W mutations are located. Yet these mutations generate substantial conformational effects within the PTC (**Supplementary Movies 1 & 2**). These long-range allosteric changes are driven through a specific network of small, yet additive, conformational changes that span across multiple domains of SBDS. A key set of interactions in domain II of WT-SBDS are the side chain interactions of Arg126 with Glu123 which in turn positions Asn121 in close proximity to Tyr128 (**Fig. 8A; Supplementary Fig. 18A**). However, the R126T substitution disrupts both these key interactions (**Fig. 8B; Supplementary Fig. 18B**). Intriguingly, even the R175W substitution perturbs this network of interactions (**Fig. 8C; Supplementary Fig. 18C**). Given that the R175W substitution resides in domain III, these changes must be driven through long-range allostery propagated within SBDS.

In SBDS-R175W, these long-range conformational changes stem from a new network of hydrophobic interactions in domain III that are driven by the mutation. The folding of Domain III arises from a network of hydrophobic interactions promoted by Leu174, Phe176, Ile177, Leu178, Ile208, Val236 and Leu234 (**Fig. 9A**). However, presence of the lone charged Arg175 residue in this hydrophobic patch in both SBDS-WT (**Fig. 9A**) and SBDS-R126T (**Fig. 9B**) allows a slight overall expansion of this domain. In the case of SBDS-R175W, replacement of the charged Arg with Trp compacts this hydrophobic patch (**Fig. 9C**) that in turn creates a series of side chain alterations across the entire length of SBDS (**Supplementary Movies 1 & 2**).

**Figure 9.**
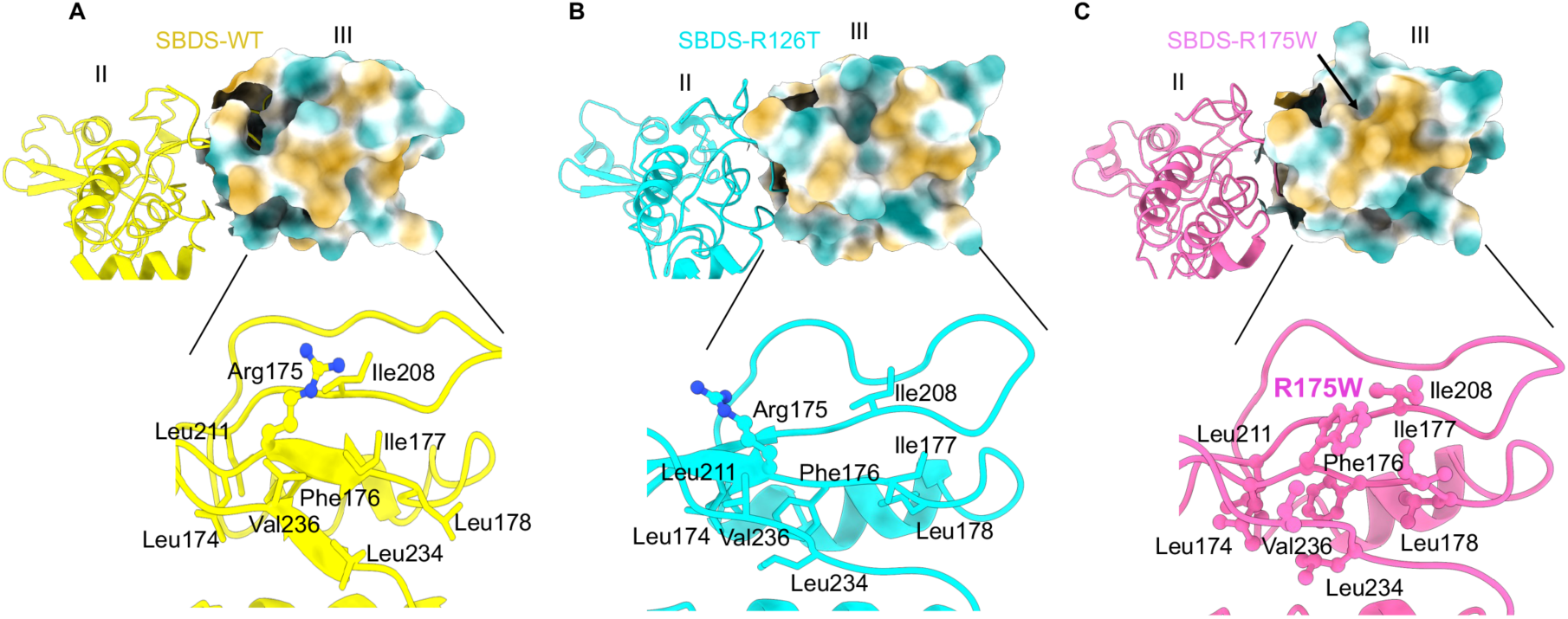
The R175W mutation remodels a hydrophobic core within SBDS domain III. *(A)* In the SBDS-WT structure, Arg175 is embedded within a hydrophobic cluster formed by Leu174, Phe176, Ile177, Leu178, Ile208, Leu234, and Val236. The presence of the positively charged Arg175 side chain interrupts this hydrophobic patch and is associated with a relatively expanded domain III conformation. *(B)* The SBDS-R126T structure retains the domain III hydrophobic network observed in SBDS-WT, including the presence of Arg175. Nevertheless, Arg175 and adjacent secondary structural elements adopt altered conformations relative to SBDS-WT, indicating that the R126T substitution in domain II propagates long-range allosteric changes into domain III. *(C)* In the SBDS-R175W structure, replacement of Arg175 with Trp introduces an additional hydrophobic residue into the core of domain III, resulting in compaction of the hydrophobic patch.

The allosteric effects of the R126T and R175W mutations observed in our structures also lead to novel interactions between domain II and the C-terminal tail of SBDS. We do not observe density for the terminal seven amino acids past Glu243 in the C-tail of SBDS-WT (**Fig. 10A; Supplementary Fig. 19A**). This finding is consistent with the disordered nature of this region in the solution NMR structure of human SBDS^9,44^. However, for both SBDS mutants, residues Glu243 through Glu247 in the C-tail forms stable interactions through a salt bridge network facilitated by Glu243, Arg169 and Glu247 (**Figs. 10B & C; Supplementary Figs. 19B & C**). For SBDS-R175W, an additional salt bridge is observed between Glu247 and Lys151. More importantly, this stable network of interactions observed in SBDS-R175W in turn stabilizes helix α5 spanning residues Asp97 to Asp117 (Domain II; **Fig. 10C**). The secondary structure assignment for this region remains ambiguous, with one solution-NMR study reporting a helical conformation and another describing the region as disordered^9,43,44^. These structures that were solved in the absence of the 60S subunit suggest that this region in SBDS may transition between ordered and disordered states. In our structures, helix α5 in SBDS-WT displays helical propensity, but the density does not fit to a strict helix (**Supplementary Fig. 20A**). In the SBDS-R126T structure, this region is disordered (**Supplementary Fig. 20B**). However, a well-ordered helix is observed for SBDS-R175W (**Supplementary Fig. 20C**). This flexible helix within domain II has been proposed to serve as a hinge region that imparts conformational flexibility to the SBDS domains, thereby enabling domain I to monitor and assess distinct conformational states. Structural transitions of this helix is an integral part of the long-range allostery in SBDS (**Supplementary Movies 1 & 2**).

**Figure 10.**
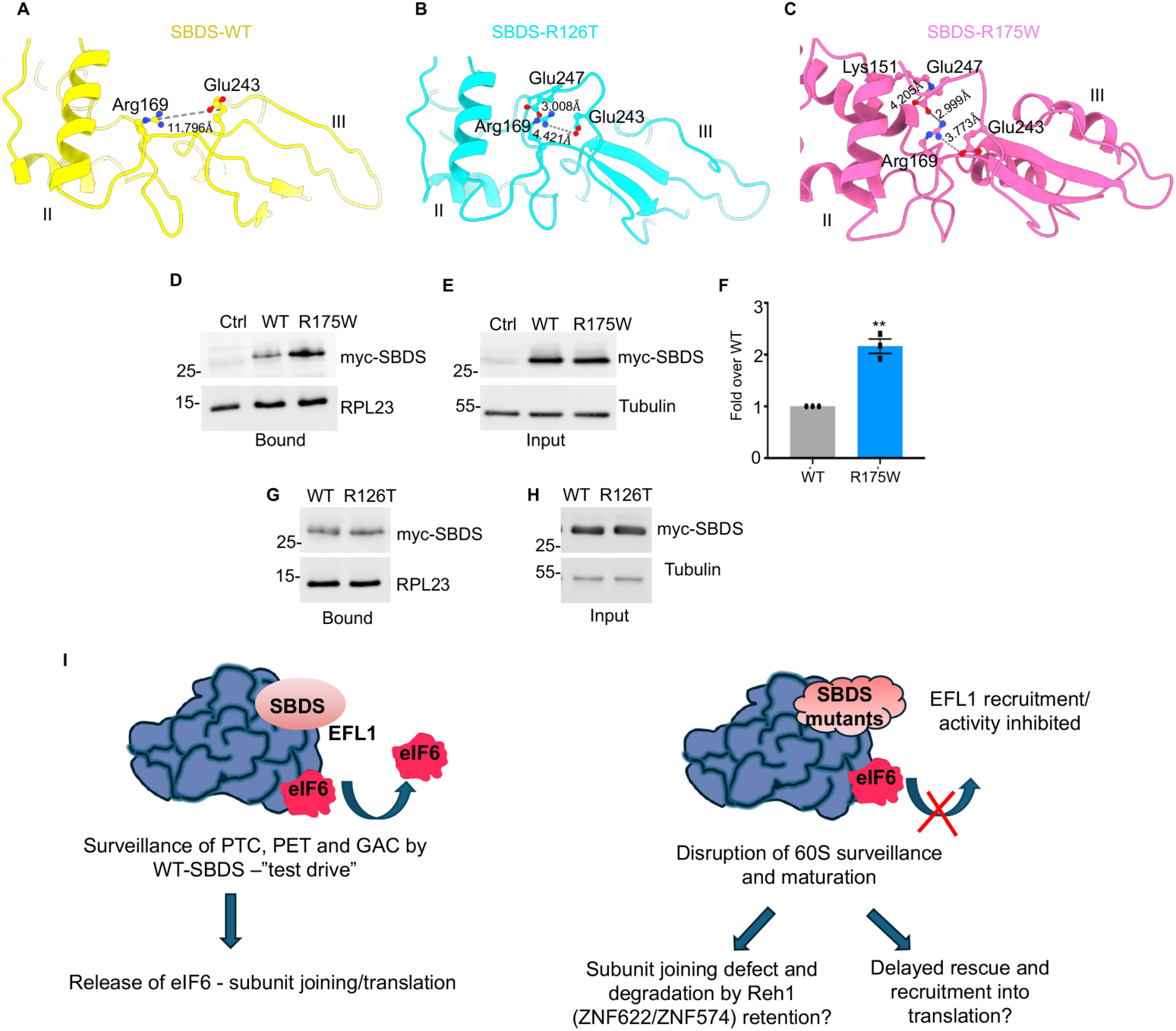
Long-range allosteric effects of SBDS mutations induce ordering of the C-terminal tail. *(A)* In the SBDS-WT structure, density for the terminal seven residues beyond Glu243 is not observed, consistent with a disordered C-terminal tail. Arg169 and Glu243 are not closely positioned. *(B)* In the SBDS-R126T structure, residues Glu243-Glu247 of the C-terminal tail are resolved and engage domain II through a salt-bridge network involving Glu243, Arg169, and Glu247. *(C)* In the SBDS-R175W structure, residues Glu243-Glu247 similarly adopt an ordered conformation stabilized by interactions with domain II through a salt-bridge network involving Glu243, Arg169, and Glu247. An additional salt bridge is observed between Glu247 and Lys151. *(D-H)* Representative western blot analysis of 60S co-sedimentation assay with transiently expressing (24 hours) myc-tagged (c-terminal) SBDS-WT, SBDS-R126T, and SBDS-R175W or empty vector control in HCT116 cells. Levels of 60S-bound and total input for WT and R175W *(D-E)* are shown. *(F)* Quantification of 60S-bound SBDS. SBDS-R175W exhibits enhanced association with the 60S subunit compared to SBDS-WT. Data represent the mean of three independent experiments and plotted as fold over WT. Significance = standard error of the mean, unpaired two-tailed *t*-Test with *p*=0.0011. *(G-H)* Similar levels of bound and total input for WT and R126T are shown. *(I)* Model on the left depicts the successful interrogation of functional centers in 60S by SBDS that triggers eIF6 release through recruitment and activation of the EFL-GTPase. Dissociation of eIF6 permits subunit joining and translation. Model on the right depicts the disruption of 60S surveillance and maturation by binding of disease variants of SBDS associated with SDS that inhibit the recruitment and/or activity of EFL1-GTPase. It is likely that these defective 60S subunits are targeted for degradation through a Reh1 (yeast) or ZNF622 or ZNF574 (human)-dependent degradation mechanism or are eventually rescued and recruited into translation by SBDS/eIF6 release.

### His101 functions as an allosteric gate for PTC interrogation

Lys98 and His101 cap one end of domain II that connects to domain 1, whereas Asp117 and Lys118 cap the other end that is connected to domain III (**Supplementary Fig. 20**). This dynamic region (domain II) that undergoes helical to disordered transitions is situated at the heart of the long-range allosteric relay in SBDS. In SBDS-R175W, Lys98 interacts with A4503 and C3791 RNA bases that support the PTC (**Supplementary Fig. 21**). This interaction is facilitated by an 180° rotation of His101, which in SBDS-WT is slotted in as a gate to block Lys98 from stably interacting with the bases in the PTC (**Supplementary Fig. 21**). Therefore, our data suggests that domain II serves as a central axle in the allosteric relay connecting spatially separated events in the P-site and the GAC (Supplementary **Fig. 20**).

A comparison of our SBDS structures reveals that the SBDS-R175W-60S complex displays the highest resolution for the three domains (**Supplementary Fig. 20**). This is due to the extensive salt-bridge and hydrogen-bonding networks observed across the domains within SBDS-R175W. Whether this stability promotes a higher binding affinity of SBDS-R175W to 60S has not been assessed in a cellular setting. We therefore transiently expressed SBDS-WT, SBDS-R126T, or SBDS-R175W-SBDS in human cells to determine their efficiency of binding to 60S. In agreement with our structural data, the SBDS-R175W mutant showed much stronger association with 60S (greater than 2-fold) compared to SBDS-WT (**Figs. 10D-F**). In contrast, SBDS-R126T bound to 60S with similar efficiency as SBDS-WT as previously reported^16^. These data suggest that SBDS-R175W could function as a dominant negative regulator to inhibit efficient release of SBDS.

## Discussion

Our findings reveal that the SDS-associated mutations of SBDS lead to defects in 60S surveillance and maturation despite distinct structural perturbations. The key contacts between SBDS and the 60S subunit are established through insertion of the N-terminal residues of SBDS into the P-site, where they engage the P-site loop of uL16 to form a mutually stabilizing interaction. Correspondingly, mutations within the uL16 P-site loop have been shown to diminish SBDS binding, underscoring the reciprocal dependence of SBDS and uL16 for stable occupancy of the P-site^18,20^. These observations support the model in which SBDS acts as a late-stage surveillance factor that monitors the integrity of critical functional centers in the 60S subunit including the P-site, peptidyl transferase center, peptide exit tunnel entrance, and GTPase-associated center. By establishing a network of interactions across these spatially distinct sites on 60S, SBDS acts as a licensing agent that assesses the structural integrity of the nascent 60S subunit and couples successful surveillance with eIF6 eviction. The two disease-variants disrupt disparate nodes within this surveillance network, providing a mechanistic explanation for how diverse structural defects yield a common failure in 60S surveillance and eIF6-eviction (**Supplementary movies 1 & 2**).

The marked but distinct structural effect of each mutant underscores the compound heterozygosity associated with the disease, wherein one mutant partly compensates for the 60S dysfunction imparted by the other mutant and vice versa. Intriguingly, there are relatively fewer disease-associated mutations in domain III of SBDS than in domains I and II^9^. Given the pronounced structural effects of the R175W mutation on domain III and its proximity to the GTPase activation center, it is possible that mutations in this region are particularly deleterious, potentially acting through dominant-negative mechanisms. We also observe distinct behavior of SBDS-R175W in solution compared to the ribosome-bound state. Although the SBDS-R175W mutant exhibits reduced thermal stability in solution (**Supplementary Fig. 1**), it is the most ordered of the ribosome-bound SBDS complexes in our structures (**Fig. 1**). These findings suggest that the R175W substitution destabilizes free SBDS while stabilizing a 60S-bound conformation, potentially contributing to the enhanced association with pre-60S ribosomal subunits observed in our pull-down experiments (**Fig. 10**). More broadly, these results underscore the importance of studying patient-associated SBDS variants in the context of the 60S-bound state whenever possible.

Structurally, SBDS has been proposed to be a tRNA mimic that test-drives the 60S subunit for functionality^9^. Successful surveillance by SBDS permits subunit joining through the EFL1-mediated eviction of eIF6, followed by the release of both SBDS and EFL1^8,9^. However, the timing and mechanism of SBDS release and whether it involves the GTPase activity of EFL1 remains unclear. The R126T substitution in SBDS has been shown to impair the 60S-dependent stimulation of EFL1 GTPase activity^9^. While the molecular basis of this defect remains unknown, our structure of the SBDS-R126T-60S complex provides important mechanistic insights. In our SBDS-R126T-bound 60 structure, both eL40 and the upper region of helix 89 exhibit increased conformational flexibility (**Supplementary Fig. 10**). eL40 is incorporated in the late-stage cytoplasmic maturation of 60S^69^. Because eL40 and helix 89 are critical for GTPase binding and activation, these observations suggest that the R126T mutation inhibits EFL1 GTPase activity by destabilizing eL40.

Interestingly, both the R126T and R175W mutants exhibit reduced binding to EFL1 in solution^49,50^, suggesting that mutation-induced alterations in SBDS structure and/or interdomain dynamics may impair EFL1 recruitment to the 60S subunit. In the case of R175W, we observe stabilization of the P-stalk base, a region whose conformational flexibility is important for the recruitment and activation of translational GTPases^60,61,64,70–74^. Our observations of enhanced association of the R175W mutant with the 60S subunit, together with the reduced mobility of the P-stalk base, suggests that the dynamic rearrangements required for productive EFL1 recruitment and binding may be compromised. Furthermore, current models of eIF6 release propose that domain V of EFL1 competes with domain III of SBDS for binding at the P-stalk base^8^. Upon engagement of the 60S subunit, EFL1 is thought to anchor at the GTPase center by displacing SBDS domain III through an approximately 180° rotation away from the P-stalk base^8^. This rearrangement permits EFL1 to adopt an accommodated conformation that sterically overlaps with the eIF6-binding site, ultimately promoting eIF6 release^8^. The compact domain III conformation observed in our SBDS-R175W-60S complex, together with its enhanced affinity for 60S, may hinder displacement of SBDS by EFL1 and thereby prevent EFL1 binding and accommodation on the 60S. Consequently, this defect would impair eIF6 release.

The GTPase activity of EFL1 has been proposed to promote the release of SBDS from the 60S subunit^8^. However, if EFL1 GTPase activity is impaired by disease-associated SBDS mutations, as observed in SDS, it remains unclear if and when these SBDS variants would dissociate from the 60S. The fate of 60S subunits bound to the pathogenic SBDS variants also remains unknown. Notably, SDS cells exhibit defects in ribosomal subunit joining in addition to reduced steady-state levels of 60S subunits relative to healthy controls^16^, suggesting that a fraction of immature 60S particles may be targeted for degradation. We therefore speculate that 60S subunits bound by SBDS-R126T or SBDS-R175W may be subjected to a quality-control pathway analogous to the Reh1-dependent surveillance mechanism recently described in yeast^75,76^. Alternatively, these mutant SBDS proteins along with eIF6 may eventually be released by EFL1, albeit with reduced efficiency, thereby permitting entry of the 60S subunits into the translating pool. Nevertheless, given the pathogenic nature of these mutations, it is likely that dissociation of mutant SBDS will compromise the 60S surveillance mechanism and would not fully restore 60S function.

SDS-associated mutations most commonly consist of truncating mutations predicted to eliminate domains II and III, together with missense variants. Although domain I contains key ribosome-binding determinants, previous studies have demonstrated that domain I alone is insufficient for stable 60S association^8^, indicating that domains II and III are required to properly anchor and orient SBDS on the ribosome. This puts forth interesting questions on how the domain I-truncations mutations of SBDS that do not bind to 60S, perturb 60S maturation. It is likely that in the absence of SBDS surveillance, faulty 60S subunits are either degraded or enter translation but do not efficiently engage in translation.

The regions of 60S bound by SBDS are critical for ribosomal function and are protected throughout key stages of 60S maturation. However, it is possible, that in the absence of SBDS either by mutations in SBDS or by inhibition of SBDS binding through mutations in uL16, it leads to re-binding of late-stage maturation factors such as NMD3 or some stress-associated factors, which could preserve 60S in an immature state. Also, somatic mutations in eIF6 such as the N106S mutation that weakens eIF6 interaction with the 60S subunit have been shown to be beneficial for recruitment of 60S subunits into translation in SDS cells^77,78^. However, it is yet to be determined if eIF6 mutations are beneficial for restoring 60S functionality in the context of 60S-binding SBDS mutants. Recent studies in yeast have identified SNOR, a factor structurally similar to domain I of SBDS that binds to the PTC during conditions of glucose deprivation^79^. However, it is likely that SNOR and other factors with structural resemblance to domain I of SBDS have evolved to regulate stress-critical translation.

Our study provides structural insights into how mutations in SBDS impair overall 60S activity and maturation. However, the timing of SBDS and eIF6 dissociation from 60S remains to be established. Premature release of eIF6 and SBDS in the absence of 40S would lead to a futile cycle of association and dissociation from 60S. Therefore, it is possible that SBDS remains anchored in the P-site until 40S subunit association. Recent studies in yeast indicate that Reh1 is situated in the lumen of the peptide exit tunnel and the C-terminus of Reh1 closely contacts the N-terminus of SBDS at the exit tunnel entrance^59,75^. Reh1 is also situated in proximity to eL24 and eIF6, and it is likely that release of eIF6 involves a Reh1-mediated allosteric communication of surveillance between SBDS and Reh1, and in collaboration with EFL1. The human ortholog of Reh1 remains unclear, but two new factors in humans ZNF622 and ZNF574 are potential candidates. ZNF622 is important for late-stage 60S maturation and functions analogous to Reh1^80^, while ZNF574 acts as a quality control factor^81^. Future efforts will be focused on capturing snapshots of these transient 60S-bound complexes in their native state towards generating a better understanding of the mechanism of eIF6 and SBDS release.

## Methods

### Purification of SBDS

Codon optimized full-length human SBDS-WT was cloned into pRSF-DUET-1 vector and variants were generated by site-directed mutagenesis (Genscript) as described^82^. Recombinant SBDS-WT and variants with 6X-His tag at C-terminus were expressed in *E. coli* BL21(DE3) cells and purified as described^82^. Cells were cultured in 500 mL LB media supplemented with 50 μg/ml kanamycin at 37°C. When the culture reached optical density (OD) 600 of 0.4 to 0.5, cells were induced with 0.4 mM Isopropyl β-D-1-thiogalactopyranoside (IPTG) followed by shaking at 250 rpm for 20 h at 16°C. Cells were harvested by centrifugation at 3724xg for 20 min. Cells were resuspended in 40 ml of lysis buffer (50 mM Tris–HCl pH 8.0, 300 mM NaCl, 10 mM imidazole, 1 mM β-mercaptoethanol, 1mM phenylmethylsulfonyl fluoride (PMSF) and 10% glycerol) supplemented with 5X protease inhibitor cocktail (PIC) (Sigma-P2714), 2 μg/mL DNase, 20 μg/mL RNase and 1 mg/mL lysozyme. Cells were lysed by stirring for 30 min at 4°C, followed by two flash freeze-thaw cycles using liquid N_2_. The freeze-thaw cycles were followed by Dounce homogenization at 4°C. The lysate was clarified at 34957xg (rotor ID: JA 25.50) for 1h at 4°C. All subsequent purification steps were performed at 4°C. The clarified lysate was applied onto a Ni^2+^-nitrolotriacetic acid (NTA) column (gravity flow). Column was washed once with 20X bed volume (BV) of lysis buffer followed by two washes with 2X BV of wash buffer (50 mM Tris pH 8.0, 1 M NaCl, 60 mM imidazole, 1 mM β-mercaptoethanol, 1× PMSF, 3X PIC and 10% glycerol). Protein was eluted using 5X BV of elution buffer (50 mM Tris pH 8.0, 300 mM NaCl, 300 mM imidazole, 1 mM β-mercaptoethanol, 1× PMSF, 5× PIC and 10% glycerol). The eluate was collected as five equal fractions and assessed by SDS-PAGE. Fractions containing SBDS were pooled, diluted 6-fold using H0 buffer (50 mM Tris-Cl pH 8.0, 1mM β-mercaptoethanol and 10% glycerol). A 5 mL HiTrap Heparin column (Cytiva Inc.) was equilibrated with 6X column volume (CV) of H50 buffer (50 mM Tris–Cl pH 8.0, 1 mM β-mercaptoethanol, 50 mM NaCl and 10% glycerol). The diluted protein was loaded onto the column using an AKTA FPLC system (3ml/min flow rate). The column was equilibrated with H50 buffer (50 mM Tris-Cl pH 8.0, 1 mM β-mercaptoethanol, 50 mM NaCl and 10% glycerol), and bound proteins were eluted using a NaCl gradient from 50 mM to 1 M. Fractions corresponding to the major absorbance peak were collected and analyzed by SDS-PAGE. Fractions containing SBDS were pooled and concentrated using an Amicon Ultra 10 kDa MCWO centrifugal concentrator and centrifuged at 4,000 rpm for 10 min. Concentrated protein was dialyzed overnight in storage buffer (50 mM HEPES pH 7.5, 200 mM NaCl, 5 mM β-mercaptoethanol, and 10% glycerol). Small aliquots were flash frozen in liquid N2 and stored at −80°C. SBDS mutant proteins (K33E, R126T, and R175W) were generated using the NEB site-directed mutagenesis kit and were purified using the same procedure as wild-type SBDS. SBDS concentration was measured using ɛ280 11 460 M−1 cm−1.

### Purification of human 60S ribosomal subunits

60S ribosomal subunits for Cryo-EM studies were purified from adherent HeLa cells as described previously^13^. HeLa cells (ATCC) were maintained in DMEM medium (Gibco) supplemented with 10% fetal bovine serum (FBS), and 100 units/ml penicillin and 100 μg/ml streptomycin (Gibco). Cells were grown to 70–80% confluency in 15–20 of 150 mm dishes. Cells were washed twice in ice-cold phosphate buffered saline (PBS). Plates were briefly tilted to collect residual PBS, which was removed by aspiration. All further steps described below were carried out using RNase free tips and tubes, and buffers were made in nuclease-free ultra-pure water. Cells were scraped in lysis buffer (15 mM Tris–Cl pH 7.5, 300 mM NaCl, 7.5 mM MgCl_2_, 2 mM DTT, 1% Triton-X 100, 1 mg/ml heparin and 0.5× cOmplete Mini EDTA-free protease inhibitor cocktail [Roche]). Lysates were rocked at 4°C for 10 min followed by centrifugation at 10 000 rpm at 4°C for 10 min. The supernatant (∼4.5 ml) was layered onto 4 ml of ice-cold 30% sucrose cushion made just before use (20 mM Tris×Cl pH 7.5, 500 mM KCl, 30% w/v RNase free sucrose, 2 mM DTT, 10 mM MgCl_2_ and 0.1 mM EDTA pH 8.0) in centrifuge tubes (Beckman Coulter, catalog# 355630). Ribosomes were pelleted by centrifugation at 63 000g for 16–19h at 4°C using a Type 80 Ti rotor (Beckman Coulter). For all ultracentrifugation steps indicated here, acceleration was set at Max, and deceleration was set at Coast (no brake) setting (Beckman Coulter-Optima XPN 100 ultracentrifuge). Ribosome pellets were resuspended in 750 μl of resuspension buffer (20 mM Tris–Cl pH 7.5, 500 mM KCl, 7.5% w/v RNAse free sucrose, 2 mM MgCl_2_, 75 mM NH_4_Cl, 2 mM puromycin [Fisher Scientific], and 2 mM DTT) by gentle pipetting. The sticky pellet was gently scraped across with a pipette tip, followed by gentle pipetting for 10min on ice to completely dissolve the pellet and till no chunks were visible. Ribosome pellets were then incubated at 4°C for 1 h to facilitate 60S and 40S subunit separation. Solution was mixed by gentle pipetting and then incubated at 37°C for 1 h. The solution was layered onto a linear 10–30% sucrose gradient (20 mM Tris–Cl pH 7.5, 500 mM KCl, w/v RNase-free sucrose and 6 mM MgCl_2_). Step gradients were prepared by freezing individual layers in liquid nitrogen in centrifuge tubes (Beckman Coulter-catalog#326823, maximum volume of 38 ml) and gradients were stored in –80°C. Prior to use, gradients were thawed overnight at 4°C. A 6–7 mm space from the top of the tube to the sucrose layer was ensured and 750 μl of the resuspended solution was layered drop by drop onto the gradient. Gradients were centrifuged at 49,100g for 16 to 17h at 4°C in an SW32Ti rotor (Beckman). Gradients were fractionated using a Brandel (UV) gradient fractionation system and 750 μl fractions were collected at a flow rate of 1.5 ml/min. The 40S fractions (∼4 ml total) and 60S fractions (∼7.5 ml total) were pelleted by centrifugation at 63 000g for 20h at 4°C in a Type 80 Ti rotor (Beckman Coulter) as described above. The subunit pellets were resuspended in storage buffer (30 mM HEPES pH 7.3, 100 mM KCl, 2.5 mM Mg (OAc)_2_, 2 mM DTT and 6% w/v RNase-free sucrose). Subunits were diluted 1:500 or 1:1000 in nuclease-free ultrapure water and OD 260 nm was measured (Agilent Cary 60 UV/VIS spectrophotometer). Subunit concentrations were calculated as described previously such that 1 *A*_260_ unit corresponds to 25 pmol of 60S and 50 pmol of 40S respectively^13^. Subunits were aliquoted on ice in RNase-free tubes and frozen in liquid N_2_ and stored at −80°C.

### In vitro 60S-binding assay

In vitro 60S-binding assay was performed as described before^82^. SBDS was diluted to 3µM in binding buffer containing 10 mM Tris-HCl pH 7.4, 75 mM KCl, and 1.5 mM MgCl₂. All buffers used in the assay were nuclease-free. SBDS was incubated with 500 nM human 60S in a final reaction volume of 15 µL. Human 60S subunits for the binding assay were purified from HeLa cells as described before^83^ and kindly provided by Dr. Christopher Fraser (University of California, Davis). The reactions were incubated for 10 min at 30 °C, followed by the addition of 50 µL binding buffer containing 0.02% Tween-20. For the pull-down assay, 20 µL of Ni-NTA magnetic agarose bead slurry (Gold Bio) was used per reaction. The beads were washed with binding buffer and equilibrated in 20 µL of binding buffer containing 0.02% Tween-20. The reactions were then added to the equilibrated beads and gently rocked for 10 min at room temperature. The beads were collected using a magnetic stand and washed three times with binding buffer containing 0.02% Tween-20. During the first wash, approximately 50 µL of binding buffer was added to the tube cap to recover any residual reaction volume, which was then gently pipetted back into the tube. Following the final wash, the beads were resuspended in 15 µL of binding buffer and mixed with 15 µL of 2X Laemmli sample buffer containing β-mercaptoethanol. Samples were boiled for 10 min, briefly centrifuged, and equal volumes were analyzed by SDS-PAGE followed by western blotting.

### Circular Dichroism

Secondary structure analysis of SBDS proteins was performed using circular dichroism (CD) spectroscopy on a J-1000 series Circular Dichroism Spectrophotometer (Jasco) as described before^13^. Measurements were carried out at 20 °C using a nitrogen-flushed setup with a 10-mm path-length cuvette. CD spectra were collected from 200 to 260 nm and corrected against a background spectrum obtained using filtered CD reaction buffer containing 100 mM NaF, 1 mM TCEP-HCl, and 5 mM Tris-HCl pH 7.5. All SBDS proteins were initially diluted to 30 µM in SBDS storage buffer (50 mM HEPES pH 7.5, 200 mM NaCl, 1 mM β-mercaptoethanol, and 10% glycerol) and then diluted into 200 µL of CD reaction buffer to a final concentration of 3 µM. For each measurement, 10 scans were collected and averaged using a 1-nm step size and 1-nm bandwidth. For thermal melt curve, variable-temperature CD spectra were determined by monitoring the molar ellipticity at 209 nm from 20 to 80 °C in 5 °C increments. The resulting thermal melting curves were fitted to a two-state unfolding model to determine the melting temperature (Tm).

### Cell culture, transfections and sucrose sedimentation

The human colorectal carcinoma line (HCT116) was obtained from ATCC and maintained in McCoy’s 5A (modified) medium with 10% fetal bovine serum (FBS) and 100 units/ml penicillin and 100 mg/ml streptomycin (Gibco) (Penn/Strep) as described before^13^. Cells were routinely tested for absence of mycoplasma using the universal mycoplasma detection kit (ATCC). Human WT-SBDS and variants carrying a disordered myc-tag at the C-terminus were cloned into pCDNA3.1 vector (Genscript). For transfections, 2×10^6^ cells were seeded per 60 mm dish and transfected with 4µg of plasmid DNA and 20µl of Lipofectamine 2000 (Invitrogen). 24h post-transfections, cells were treated with 100 µg/ml Cycloheximide (Tocris) for 5 min at 37 °C and washed twice with cold 1X phosphate buffered saline (PBS) supplemented with 100µg/ml cycloheximide at 4°C. The cells were collected by scraping in polysome lysis buffer (PLB: 50 mM HEPES pH 7.4, 250 mM KCl, 5 mM MgCl2, 250 mM sucrose, 1% Triton X-100, 1% sodium deoxycholate, 100 μg/ml cycloheximide, 100 units/ml Superasin (Invitrogen), 0.25X protease inhibitor cocktail EDTA free (Roche, Sigma). Cells were collected in 300µl PLB per 60mm dish and two dishes were pooled per sample (600µl total). Lysates were rocked for 10min at 4°C cleared by centrifugation at 13,000 rpm for 10min. 100µL was saved as input and 500µl was layered onto 7.5 ml of 1M sucrose cushion (10 mM HEPES pH 7.4, 200 mM KCl, 10 mM MgCl_2_, 100 µg/ml cycloheximide, 1M Sucrose) in centrifuge tubes (Beckman Coulter, catalog# 355630). Ribosomes were sedimented at 70,000 rpm for 2h at 4°C using a Type 80 Ti rotor (Beckman Coulter) with slow acceleration and deceleration setting at 9 (Beckman Coulter-Optima XPN 100 ultracentrifuge). Ribosomal pellet was washed once to remove residual sucrose by gently swirling with ice-cold PLB and resuspended in 150µl of PLB and 1X Laemmli sample buffer. Samples were boiled for 7 min and same volume for each sample was probed by western blotting.

### Western blotting and antibodies

Western blotting was performed as previously described^13^. Proteins were resolved by sodium dodecyl sulfate–polyacrylamide gel electrophoresis (SDS-PAGE) and transferred onto nitrocellulose membrane (0.45 μm; Bio-Rad Laboratories). Blots were blocked for 1 h at room temperature in 5% non-fat dry milk dissolved in tris buffered saline with 0.1% Tween 20 (TBS-T). Membranes were probed overnight at 4 °C and rocked gently with the following antibodies: RPL23 (uL14) antibody (1:2500; Bethyl, A305-010A and with 5% milk for cellular assays), SBDS antibody (1:1000; Santa Cruz Biotechnology), myc-tag (1:1000 in 1% milk; Cell Signaling, 2278), α-Tubulin (1:3000; Cell signaling, 8058) and His-tag HRP (1:2500, Santa Cruz Biotechnology, sc8036). Blots were probed with the respective secondary antibodies for 1h at RT: donkey anti-rabbit whole IgG HRP (1:30000; Jackson Immunoresearch, 715-035-152) and Donkey anti-mouse whole IgG HRP (1:30000; Jackson Immunoresearch, 715-035-150). Primary and secondary antibodies were diluted in Tris-buffered saline containing 0.1% Tween 20. Blots were developed with enhanced chemiluminescence (ECL) substrate (Pierce). Images were acquired and quantitated using iBright CL1500 imager (Thermo Fisher Scientific) and iBright analysis software.

### Cryo-EM sample preparation and data collection

Cryo-EM samples were prepared with 500 nM SBDS incubated with 45 nM purified 60S in binding buffer (10 mM Tris-HCl, pH 7.4, 75 mM KCl, and 1.5 mM Mg(OAc)₂). Samples were incubated for 10 minutes at 30 °C. UltrAufoil R2/2 Au grids were negatively glow-discharged for 60 s using a GloQube (Quorum). Immediately following grid cleaning and incubation of the sample, 3 µl of the sample was applied to each grid. Grid preparation was performed using a Vitrobot Mark IV (Thermo Fisher Scientific) with the chamber maintained at 4 °C and 100% humidity. Without further incubation, grids were blotted for 2.5 s (blot force 2) and plunge-frozen into liquid ethane cooled by liquid nitrogen. Grids were stored in liquid nitrogen until data collection. All procedures were performed under RNase-free conditions. Grids were clipped and loaded into either a 200-kV Thermo Fisher Scientific Glacios or a 300-kV Thermo Fisher Scientific Krios transmission electron microscope. Data were acquired using EPU software (Thermo Fisher Scientific) with aberration-free image shift and recorded on a Falcon IV direct electron detector (4k × 4k). Although minor preferred orientation was observed, several optimization strategies, including the use of different grid types and freezing conditions, were explored but did not substantially improve the orientation distribution. Nevertheless, all datasets exhibited good particle distribution and sufficient orientation diversity, as assessed by two-dimensional classification.

### Data processing

#### Vacant-60S

Single-particle cryo-EM data were processed using cryoSPARC v.5.0.8. For the vacant 60S dataset, 9,891 raw videos were motion-corrected using patch motion correction, followed by patch contrast transfer function (CTF) estimation (**Supplementary Fig. 2**). Initial particle picking was performed using a template picker, and particles were extracted with a box size of 700 pixels (binning factor 1.5). This yielded an initial stack of 624,240 particles. Several rounds of two-dimensional classification were conducted to remove false positives and damaged particles, resulting in a cleaned subset of 154,139 particles. These particles were used to generate three classes of *ab initio* three-dimensional reconstructions. One of the 3D reconstruction corresponding to the well-defined 60S ribosome (69,540 particles) was subjected to homogeneous refinement followed by NU-refinement. Resulting final map reached a resolution of 2.49 Å at a Fourier shell correlation (FSC) threshold of 0.143.

#### SBDS-WT-60S complex

Single-particle cryo-EM data were processed using cryoSPARC v.5.0.8. For the WT-SBDS bound 60S dataset, 3,324 raw motion corrected micrographs followed by patch contrast transfer function (CTF) estimation (**Supplementary Fig. 3**). Initial particle picking was performed using a template picker, and particles were extracted with a box size of 700 pixels (binning factor 4). This yielded an initial stack of 164,585 particles. Several rounds of two-dimensional classification were conducted to remove false positives and damaged particles, resulting in a cleaned subset of 52,039 particles. These particles were used to generate initial *ab initio* three-dimensional reconstructions followed by homo and NU refinement. This particle subsets were then re-extracted to its original box size yielding a particle stack of 51,536 particles which was followed by another round of homo and NU-refinement. Density for SBDS region was further improved by applying focused mask on the SBDS region including regions around as well on the 60S and running it though local refinement job. This yielded an improved map for the SBDS region while the overall final map was further sharpened using DeepEMhancer. The overall map had a global resolution of 2.92 Å (FSC = 0.143), whereas the focused map had a resolution of 2.96 Å (FSC = 0.143).

#### SBDS-R126T-60S complex

Single-particle cryo-EM data were processed using cryoSPARC v. 5.0.8. For the R126T-SBDS-bound 60S dataset, a total of 13,859 raw movies, collected at 0° and 30° tilt, were processed separately up to the 2D classification step and cleaner classes merged together. Motion correction was performed using Patch Motion Correction, followed by contrast transfer function (CTF) estimation using Patch CTF Estimation (**Supplementary Fig. 4**). Initial particle picking was performed using a template picker, and particles were extracted using a box size of 700 pixels (binning factor 4), yielding an initial particle stack of 480,405 particles. Several rounds of two-dimensional classification were performed to remove false-positive picks and damaged particles, resulting in a cleaned subset of 123,088 particles. These particles were used to generate initial *ab initio* three-dimensional (3D) reconstructions, followed by heterogeneous refinement, non-uniform (NU) refinement, and an additional round of heterogeneous refinement using the best-resolved class. The well-defined 3D reconstruction of the SBDS-bound 60S complex was subjected to another round of NU refinement, followed by 3D classification using a mask encompassing SBDS and the surrounding region of the 60S ribosome. This classification yielded a cleaner particle stack of 76,049 particles. The corresponding 3D reconstruction was further refined through an additional round of 3D classification using a mask focused on SBDS. The selected particles were then re-extracted at the original box size and subjected to local refinement using a focused mask encompassing SBDS. This procedure improved the density corresponding to SBDS, yielding a final reconstruction at an overall resolution of 3.17 Å (FSC = 0.143), based on 18,085 particles.

#### SBDS-R175W-60S

Single-particle cryo-EM data were processed using cryoSPARC v. 5.0.8. For the R175W-SBDS-bound 60S dataset, a total of 6,168 raw movies were subjected to motion correction using Patch Motion Correction, followed by contrast transfer function (CTF) estimation using Patch CTF Estimation (**Supplementary Fig. 5**). Initial particle picking was performed using a template picker, and particles were extracted using a box size of 512 pixels (binning factor 2), yielding an initial particle stack of 485,079 particles. Several rounds of two-dimensional (2D) classification were performed to remove false-positive picks and damaged particles, resulting in a cleaned subset of 31,104 particles. These particles were used to generate two initial *ab initio* three-dimensional (3D) reconstructions, followed by 3D classification into three classes using the well-resolved class. The best-resolved class was subsequently subjected to non-uniform (NU) refinement, followed by 3D variability analysis using five classes in the 3D Variability Display, with a mask encompassing the SBDS region. This analysis was performed to improve the density of SBDS bound to the 60S ribosome. Two classes, containing 3,893 and 10,805 particles, respectively, showed well-resolved density for the SBDS region. These particle subsets were combined and subjected to a final round of NU refinement, yielding the final reconstruction with a global resolution of 3.20 Å (FSC = 0.143).

#### SBDS-K33E-60S complex

Single-particle cryo-EM data were processed using cryoSPARC v. 5.0.8. For the K33E-SBDS-bound 60S dataset, a total of 6,950 raw movies, collected at 0° and 30° tilt, were processed separately up to the 2D classification step and cleaner classes merged together.

Motion correction was performed using Patch Motion Correction, followed by contrast transfer function (CTF) estimation using Patch CTF Estimation (**Supplementary Fig. 7**). Initial particle picking was performed using a template picker, and particles were extracted using a box size of 700 pixels (binning factor 2), yielding an initial particle stack of 251,389 particles. Several rounds of two-dimensional (2D) classification were performed to remove false-positive picks and damaged particles, resulting in a cleaned subset of 100,584 particles. These particles were used to generate initial *ab initio* three-dimensional (3D) reconstructions, followed by heterogeneous refinement and non-uniform (NU) refinement of the better-resolved reconstruction containing SBDS bound to the 60S ribosome. The resulting particle set was further subjected to heterogeneous refinement, yielding three distinct volumes: one corresponding to a well resolved 60S ribosome without bound SBDS, a second showing SBDS associated with the 60S ribosome, and a third poorly resolved volume. Notably, the SBDS-bound volume exhibited a substantially different structural appearance compared with the corresponding reconstructions obtained for other SBDS mutants and the wild-type (WT) complex. This observation was reproducible across multiple independent processing attempts. To further investigate and resolve this distinct class, an additional round of heterogeneous refinement was performed; however, the resulting SBDS-bound volume remained poorly resolved.

### Model building and refinement

Previously determined structures, including PDB entries 5AN9, 6QKL and 7OW7, were used as starting models for fitting into the electron microscopy density maps of all complexes using ChimeraX. The resulting models were iteratively rebuilt and refined in Coot v.0.9.8.93, followed by further refinement and validation using PHENIX v.1.21.2-5419. Structural figures were generated using ChimeraX.

## Supporting information

All Supplemental figures

## Acknowledgements

This work was supported by funding from the National Institutes of Health - National Institute of General Medical Science R01-GM143179 (S.O.), R35-GM149320 (E.A.); the National Cancer Institute R01-CA305375 (E.A.). We thank Drs. Summer Brock, Katherine Basore, and Bradley Readnour at the Washington University Center for Cellular Imaging; Dr. Ed Eng at NCCAT for Cryo-EM data collection; and Dr. Greg Sabat, Mass Spectrometry Core at University of Wisconsin Madison. We also thank Dr. Ahmad Jomaa at the University of Virginia School of Medicine, for his technical and intellectual input.

## Code Availability Statement

No custom code was used in this manuscript.

## Data Availability Statement

All raw data are contained within the manuscript and provided in the Source Data File. Plasmids used for protein overexpression are available upon request. (The single particle Cryo-EM maps and models are being deposited into PDB and EMDB and codes will be updated soon).

## Author Contributions

Conceptualization: SO. Experimental Design: SO, and EA. Methodology & Experiments: RK (solved all structures), SA (biochemistry), PR (cellular study), SO, and EA. Visualization: RK, SO, and EA. Funding acquisition: SO, and EA. Project administration: SO, and EA. Supervision: SO, and EA. Writing-SO and EA, Methods- RK, SA and PR, Writing - review & editing: all authors.

