## Supplementary material for "Structural basis of SBDS dysfunction in Shwachman-Diamond Syndrome": All Supplemental figures

**Supplemental Movies 1-2**
**Supplemental Figures 1 – 21**

**Supplementary Movie 1. Long-range allosteric remodeling of SBDS on the 60S ribosomal subunit induced by the SDS-associated R126T mutation.** Morphing animation comparing the SBDS-WT-60S (starting model) and SBDS-R126T-60S (ending model) structures. SBDS is shown in cyan and comprises three domains (I-III), with domain II forming the central helical region that mediates long-range allosteric communication across the protein. Residues Lys98 and His101 cap one end of the central helix (hinge between domains I and II), whereas Asp117 and Lys118 cap the opposite end; these residues undergo large-scale rotational movements associated with mutation-induced conformational remodeling and the helical definition is lost due to mutation. uL16 (RPL10) is shown in blue. Rotational changes in A3908 and A4548 bases are shown as sticks and are examples of corresponding changes in the 60S that are induced by the SBDS-R126T mutation. The sarcin-ricin loop (bases 4594-4613) is also depicted.

**Supplementary Movie 2. Long-range allosteric remodeling of SBDS on the 60S ribosomal subunit induced by the SDS-associated R175W mutation.** Morphing animation comparing the SBDS-WT-60S (starting model) and SBDS-R175W-60S (ending model) structures. SBDS is shown in cyan and comprises three domains (I-III), with domain II forming the central helical region that mediates long-range allosteric communication throughout the protein. Residues Lys98 and His101 cap one end of the central helix (hinge between domains I and II), whereas Asp117 and Lys118 cap the opposite end; these residues undergo large-scale rotational movements associated with mutation-induced conformational remodeling and R175W mutation stabilizes this alpha-helix. uL16 (RPL10) is shown in blue. Rotational changes in A3908 and A4548 bases are shown as sticks and are examples of corresponding changes in the 60S that are induced by the SBDS-R175W mutation. The sarcin-ricin loop (bases 4594-4613) is also depicted.

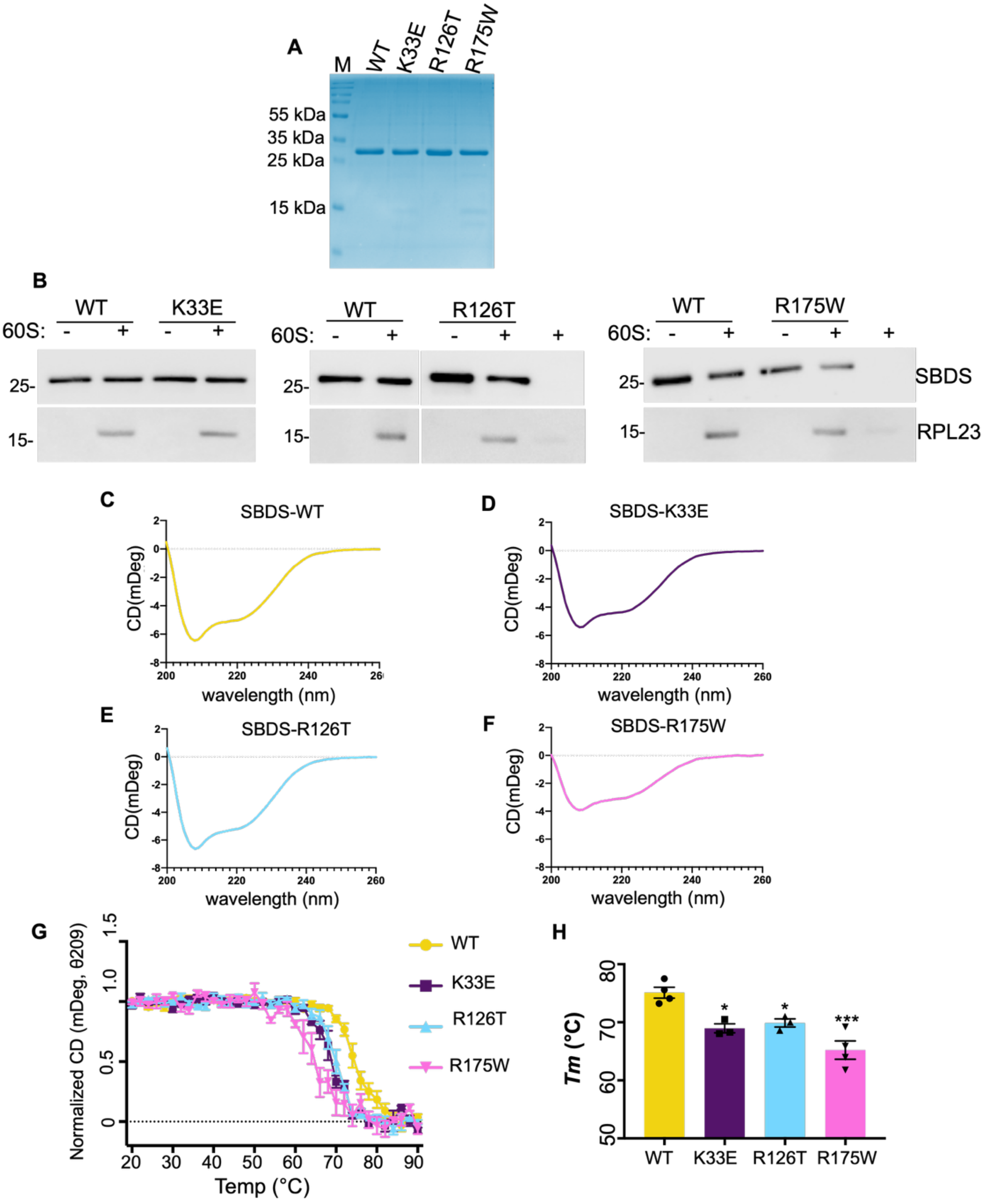


**Supplemental Figure 1. Ribosomal association and protein stability properties of SBDS mutants.** *(A)* SDS-PAGE analysis of purified recombinant SBDS-WT, SBDS-K33E, SBDS-R126T, and SBDS-R175W proteins used in this study. Data represent three-independent repeats

*(B)* Pull-down and immunoblot analysis of 60S ribosomal subunits and 60S-bound SBDS complexes comparing SBDS-WT with the indicated disease-associated variants. *(C, D, E and F)* Circular dichroism (CD) spectra of SBDS-WT and mutant proteins. All variants exhibit similar overall spectra indicating preservation of global secondary structure. However, SBDS-K33E and SBDS-R175W display reduced overall ellipticity consistent with differences in the extent of folded secondary structural elements. Data represent three-independent repeats. *(G)* Thermal denaturation analysis monitored by CD changes at 209 nm. Mean residue ellipticity (mDeg) is plotted as a function of temperature to assess protein unfolding. *(H)* Quantitation of melting temperatures ($T_m$) derived from the thermal denaturation curves. Mean $T_m$: SBDS-WT = 75.11°C, SBDS-K33E=68.96°C, SBDS-R126T = 69.88°C, and SBDS-R175W= 65.21°C. Data represent three independent repeats and standard error of the mean. Statistical significance calculated using one-way ANOVA (Dunnett's multiple comparisons test) with p=0.0105 (WT vs. K33E), p=0.0260 (WT vs. R126T) and p=0.0002 (WT vs. R175W).

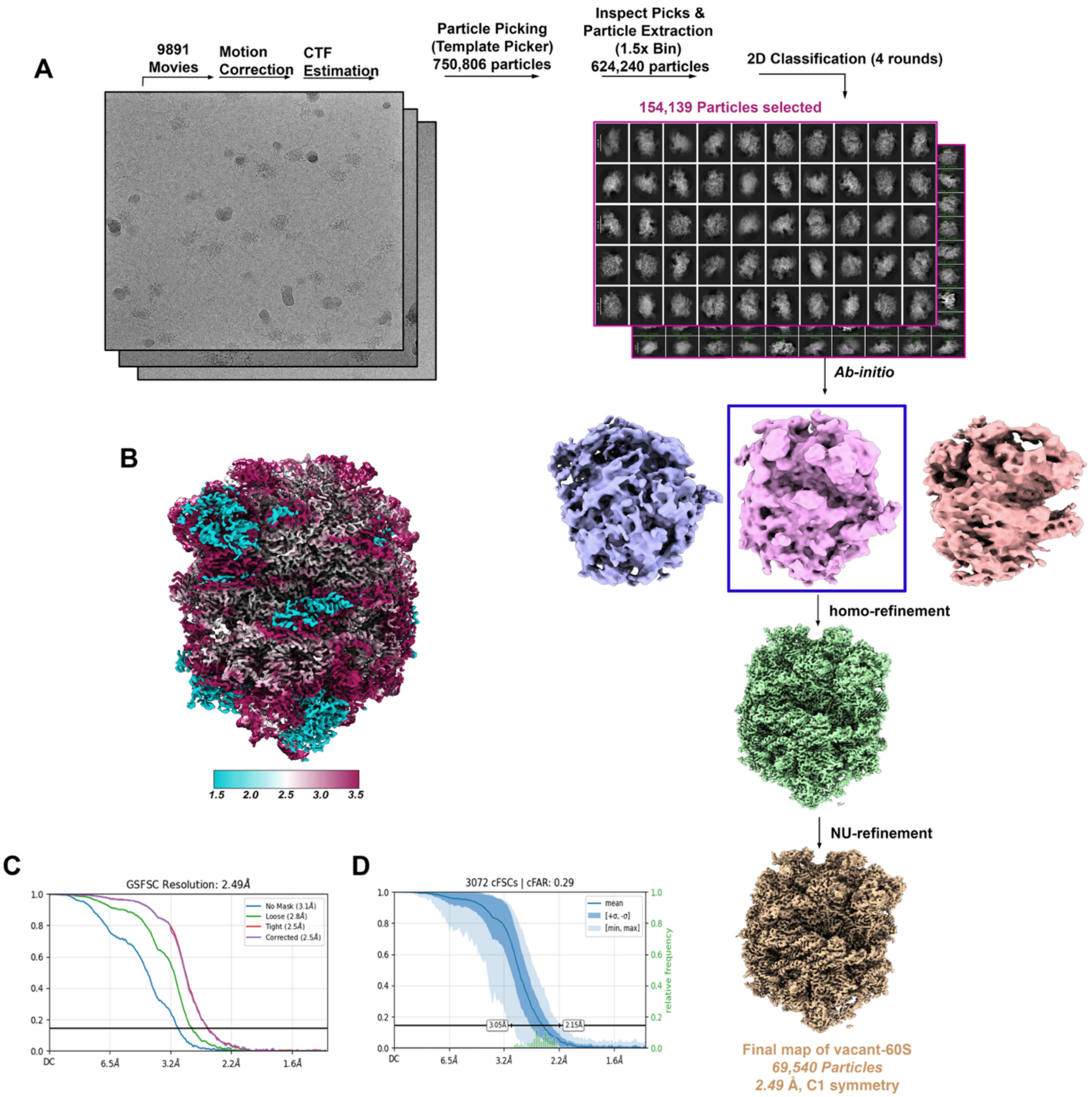


**Supplemental Figure 2. Cryo-EM data processing workflow for vacant 60S.** *(A)* Flowchart of single-particle cryo-EM image processing for the vacant 60S complex. Colored boxes indicate particle subsets and corresponding 3D reconstructions selected for further processing. *(B)* Local resolution map of the final vacant 60S reconstruction. *(C)* Gold-standard Fourier shell correlation (FSC) curve for reconstruction, showing an overall resolution of 2.49 Å, determined using the FSC = 0.143 criterion. *(D)* 3D directional Fourier shell correlation (3D-FSC) plot for the final vacant 60S reconstruction. The blue curve represents the directional FSC average, and the shaded region indicates the spread among different orientations, demonstrating isotropic resolution distribution. The dashed horizontal line marks the FSC = 0.143 threshold.

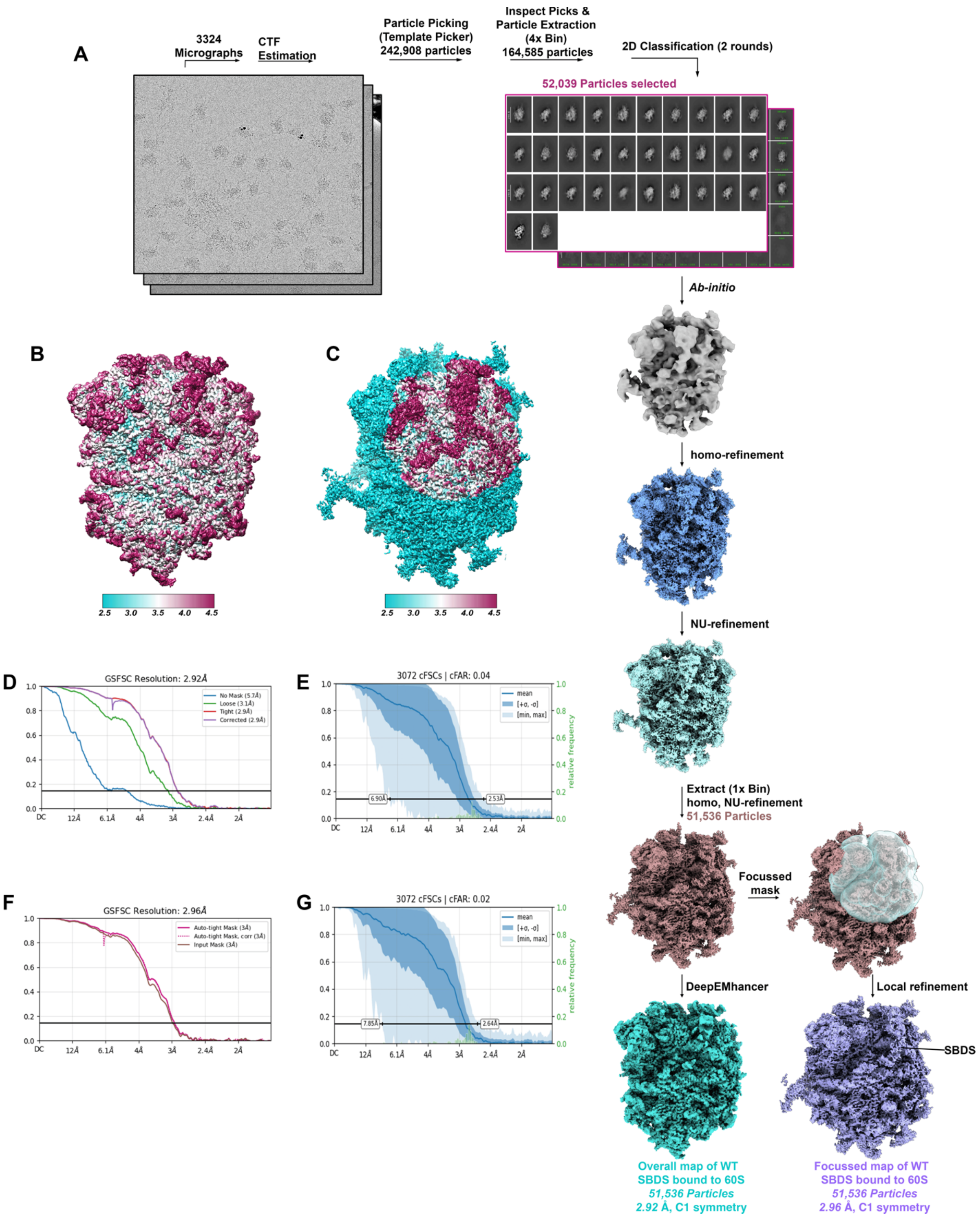


**Supplemental Figure 3. Cryo-EM data processing workflow for the SBDS-WT-60S complex.** *(A)* Flowchart of single-particle cryo-EM image processing for the SBDS-WT bound 60S complex. Colored boxes indicate particle subsets, and corresponding 3D reconstructions selected for further processing. *(B)* & *(C)* Local resolution maps of the overall map and focused map of SBDS bound

60S reconstruction, respectively. *(D)* Gold-standard Fourier shell correlation (FSC) curve for the overall reconstruction, showing an overall resolution of 2.92 Å, determined using the FSC = 0.143 criterion. *(E)* 3D directional Fourier shell correlation (3D-FSC) plot for the final SBDS-WT bound 60S complex reconstruction. *(F)* Gold-standard Fourier shell correlation (FSC) curve for the focused map reconstruction, showing an overall resolution of 2.96 Å, determined using the FSC = 0.143 criterion. *(G)* 3D directional Fourier shell correlation (3D-FSC) plot for the focused map of SBDS-WT bound 60S complex reconstruction. The blue curve represents the directional FSC average, and the shaded region indicates the spread among different orientations, demonstrating isotropic resolution distribution. The dashed horizontal line marks the FSC = 0.143 threshold.

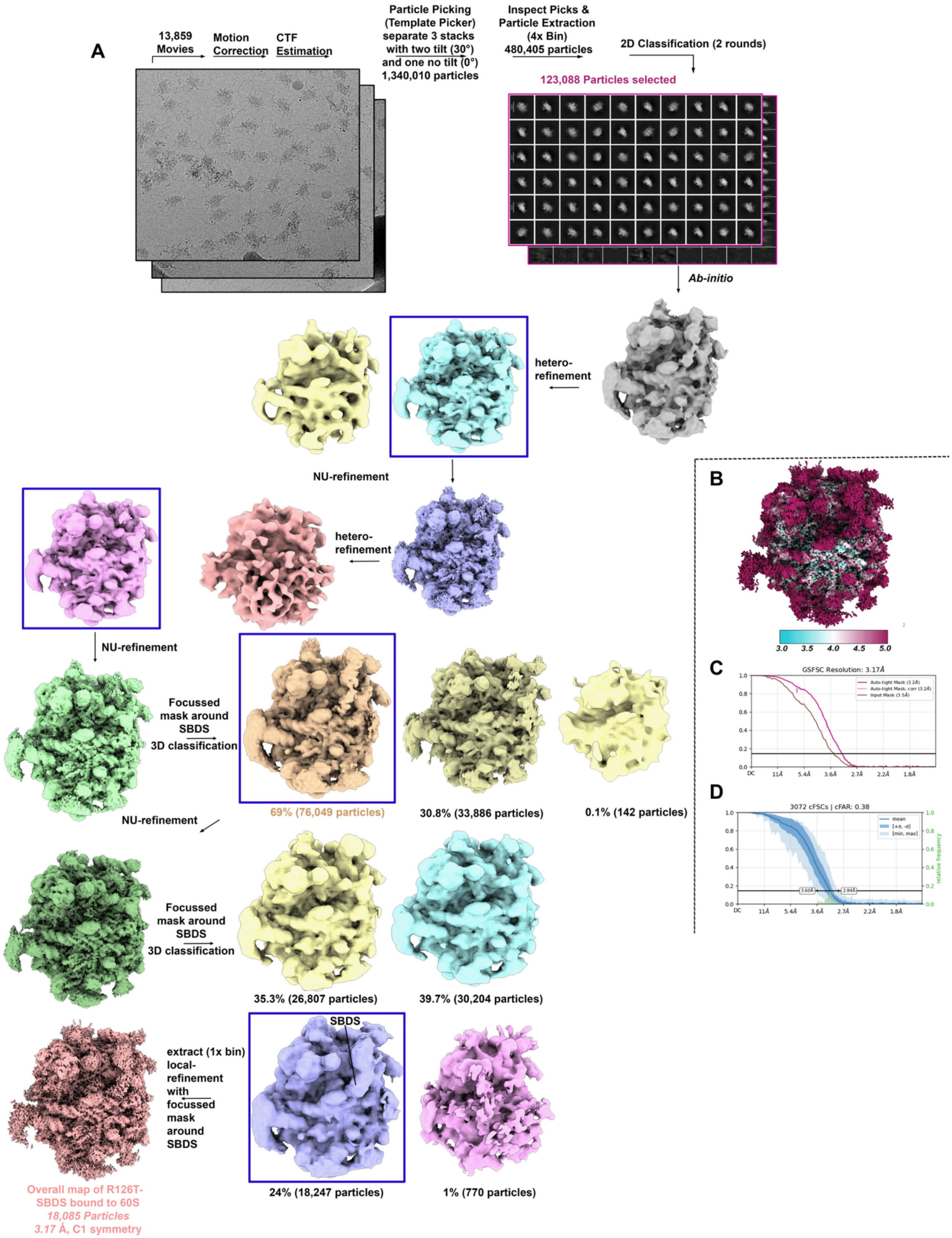
A
13,859 Movies
Motion Correction
CTF Estimation
Particle Picking (Template Picker) separate 3 stacks with two tilt (30°) and one no tilt (0°) 1,340,010 particles
Inspect Picks & Particle Extraction (4x Bin) 480,405 particles
2D Classification (2 rounds)
123,088 Particles selected
Ab-initio
hetero-refinement
NU-refinement
hetero-refinement
NU-refinement
Focussed mask around SBDS 3D classification
69% (76,049 particles)
30.8% (33,886 particles)
0.1% (142 particles)
NU-refinement
Focussed mask around SBDS 3D classification
35.3% (26,807 particles)
39.7% (30,204 particles)
extract (1x bin) local-refinement with focussed mask around SBDS
SBDS
24% (18,247 particles)
1% (770 particles)
Overall map of R126T-SBDS bound to 60S
18,085 Particles
3.17 Å, C1 symmetry
B
3.0
3.5
4.0
4.5
5.0
C
GSFSC Resolution: 3.17Å
D
3072 cFSCs | cFAR: 0.38

**Supplemental Figure 4. Cryo-EM data processing workflow for the SBDS-R126T-60S complex.** *(A)* Flowchart of single-particle cryo-EM image processing for the SBDS-R126T bound 60S complex. Colored boxes indicate particle subsets, and corresponding 3D reconstructions selected for further processing. *(B)* Local resolution map of the final SBDS-R126T bound 60S reconstruction. *(C)* Gold-standard Fourier shell correlation (FSC) curve for the reconstruction, showing an overall resolution of 3.17 Å, determined using the FSC = 0.143 criterion. *(D)* 3D directional Fourier shell correlation (3D-FSC) plot for the final SBDS-R126T bound 60S reconstruction. The blue curve represents the directional FSC average, and the shaded region indicates the spread among different orientations, demonstrating isotropic resolution distribution. The dashed horizontal line marks the FSC = 0.143 threshold.

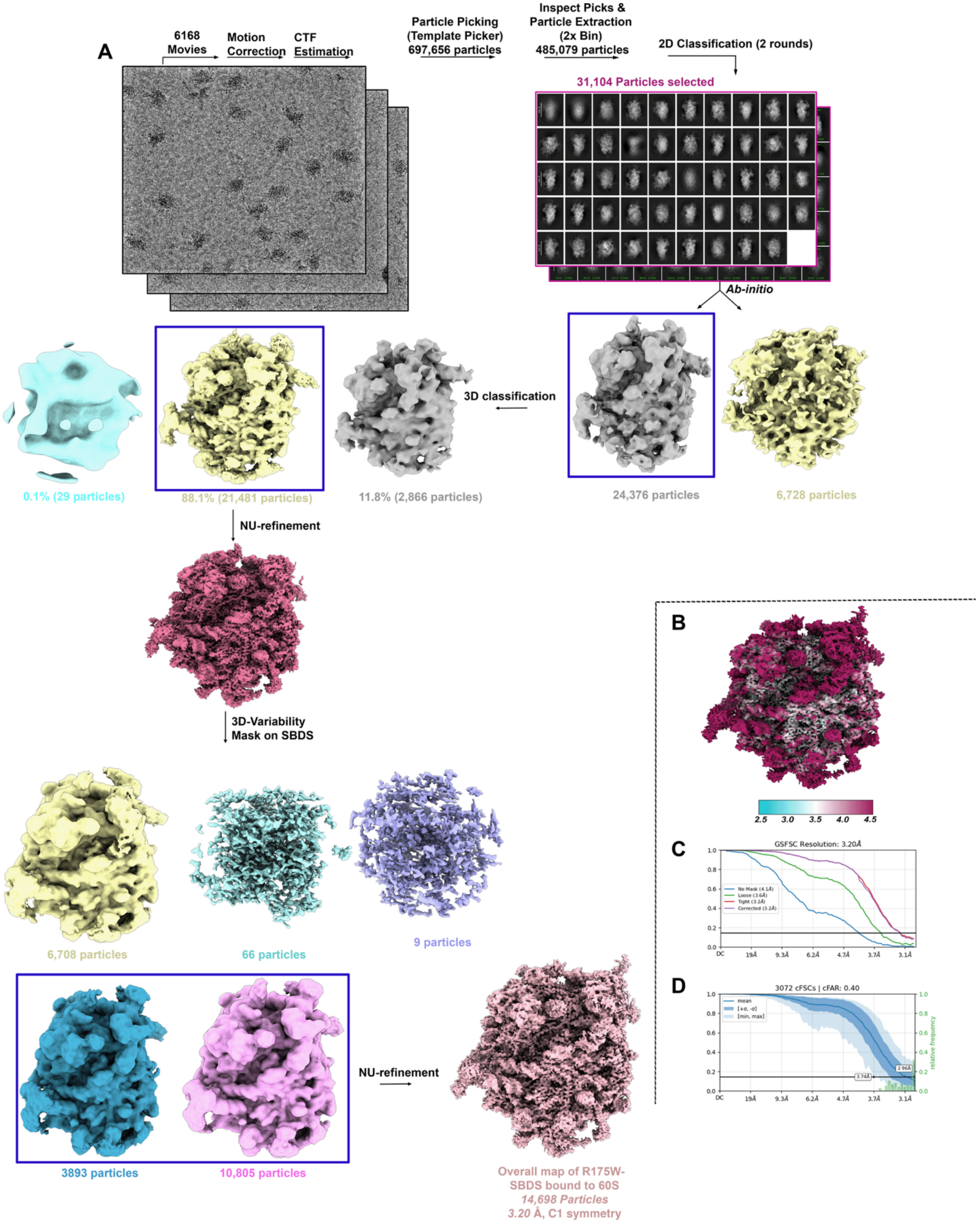


**Supplemental Figure 5. Cryo-EM data processing workflow for the SBDS-R175W-60S complex.** *(A)* Flowchart of single-particle cryo-EM image processing for the SBDS-R175W bound 60S complex. Colored boxes indicate particle subsets, and corresponding 3D reconstructions selected for further processing. *(B)* Local resolution map of the final SBDS-R175W bound 60S reconstruction. *(C)* Gold-standard Fourier shell correlation (FSC) curve for the reconstruction, showing an overall resolution of 3.20 Å, determined using the FSC = 0.143 criterion. *(D)* 3D directional Fourier shell correlation (3D-FSC) plot for the final SBDS-R175W bound 60S reconstruction. The blue curve represents the directional FSC average, and the shaded region indicates the spread among different orientations, demonstrating isotropic resolution distribution. The dashed horizontal line marks the FSC = 0.143 threshold.

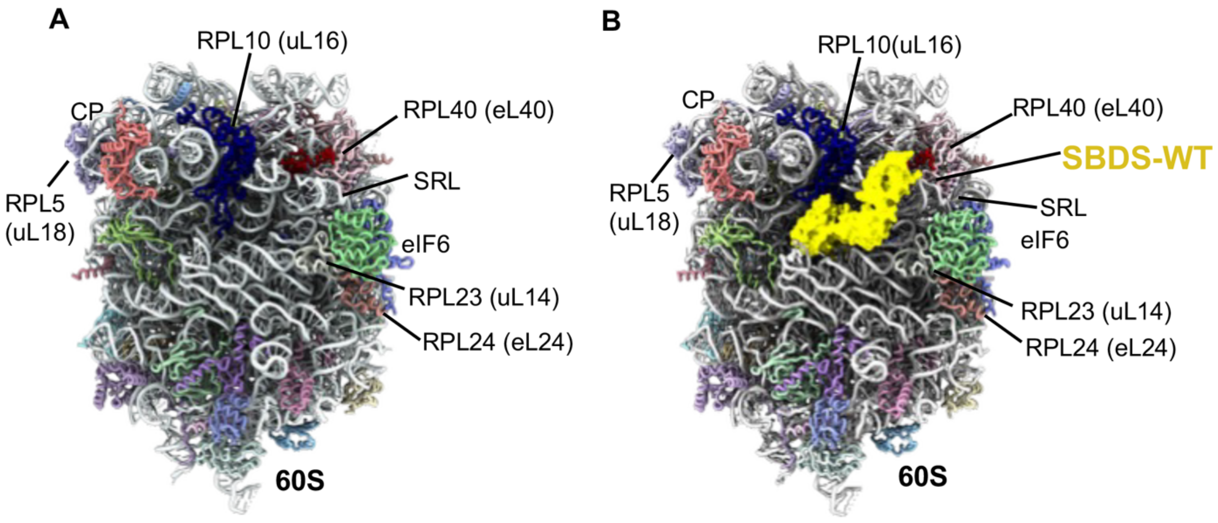


**Supplemental Figure 6. Position of eIF6 modeled on 60S relative to SBDS.** eIF6 (green) is docked onto the structure of *(A)* vacant or *(B)* SBDS-WT bound 60S. The docked model is based on PDB: 6QKL. The binding positions of SBDS and eIF6 reveal that there are no direct contacts.

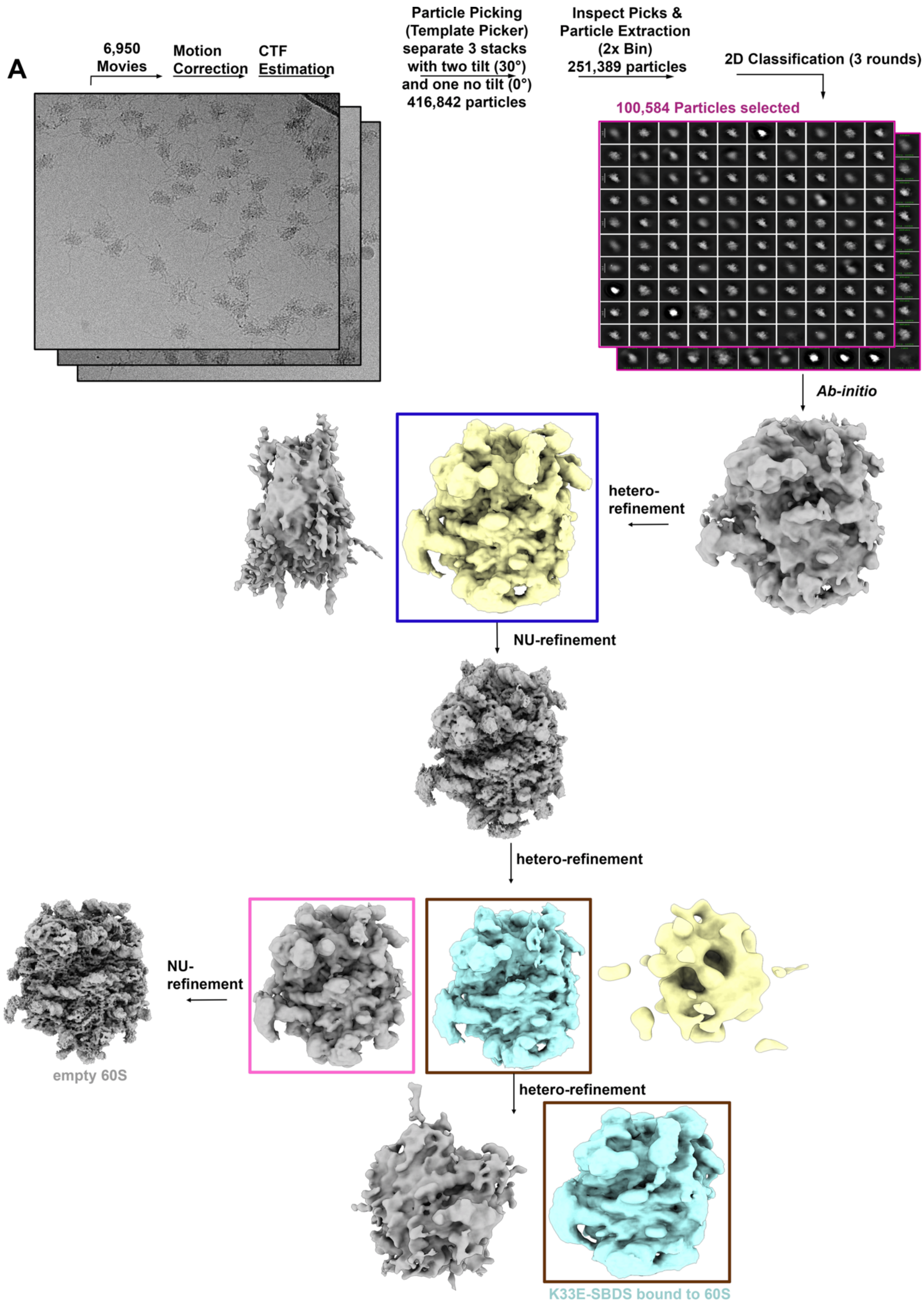


**Supplemental Figure 7. Cryo-EM data processing workflow for the SBDS-K33E-60S complex.** *(A)* Flowchart of single-particle cryo-EM image processing for the SBDS-K33E bound

60S complex. Processing from one of four distinct trials is shown. All four datasets encountered the same problems shown here. Colored boxes indicate particle subsets, and corresponding 3D reconstructions selected for further processing. The best volume of the SBDS-K33E is shown in the bottom right. The volumes were poor and not resolved further. The vacant 60S molecules from the same grid separated out and were resolvable to very high resolutions.

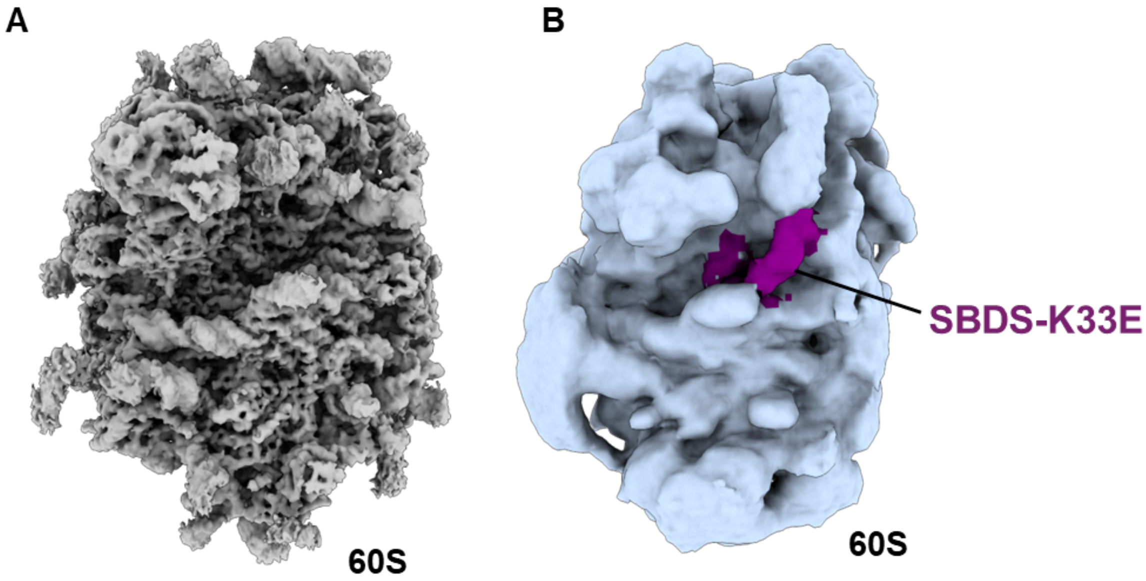


**Supplemental Figure 8. Poor volumes observed for the SBDS-K33E complex.** *(A)* Vacant 60S from the SBDS-K33E-60S grids produced volumes with high resolution (3.17Å). *(B)* A very small subset of the particles ~49000 out of ~1.3 million 60S in the cryo-EM dataset possessed density for SBDS-K33E (highlighted in purple).

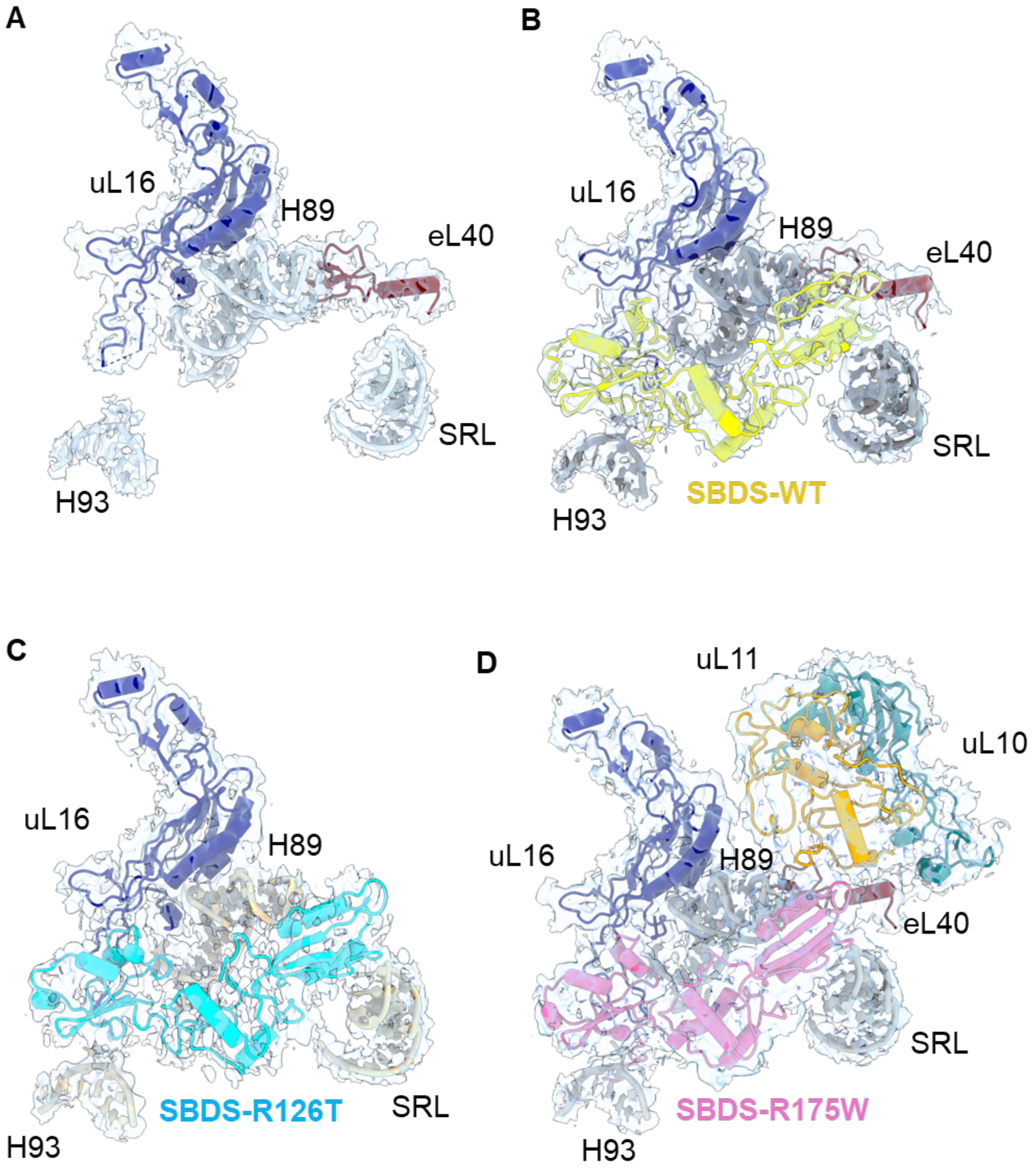


**Supplementary Figure 9. Density in the cryo-EM data.** Carved density for structures shown in Figure 2. Panels *(A-D)* here correspond to structures in Figure 2, panels A-D.

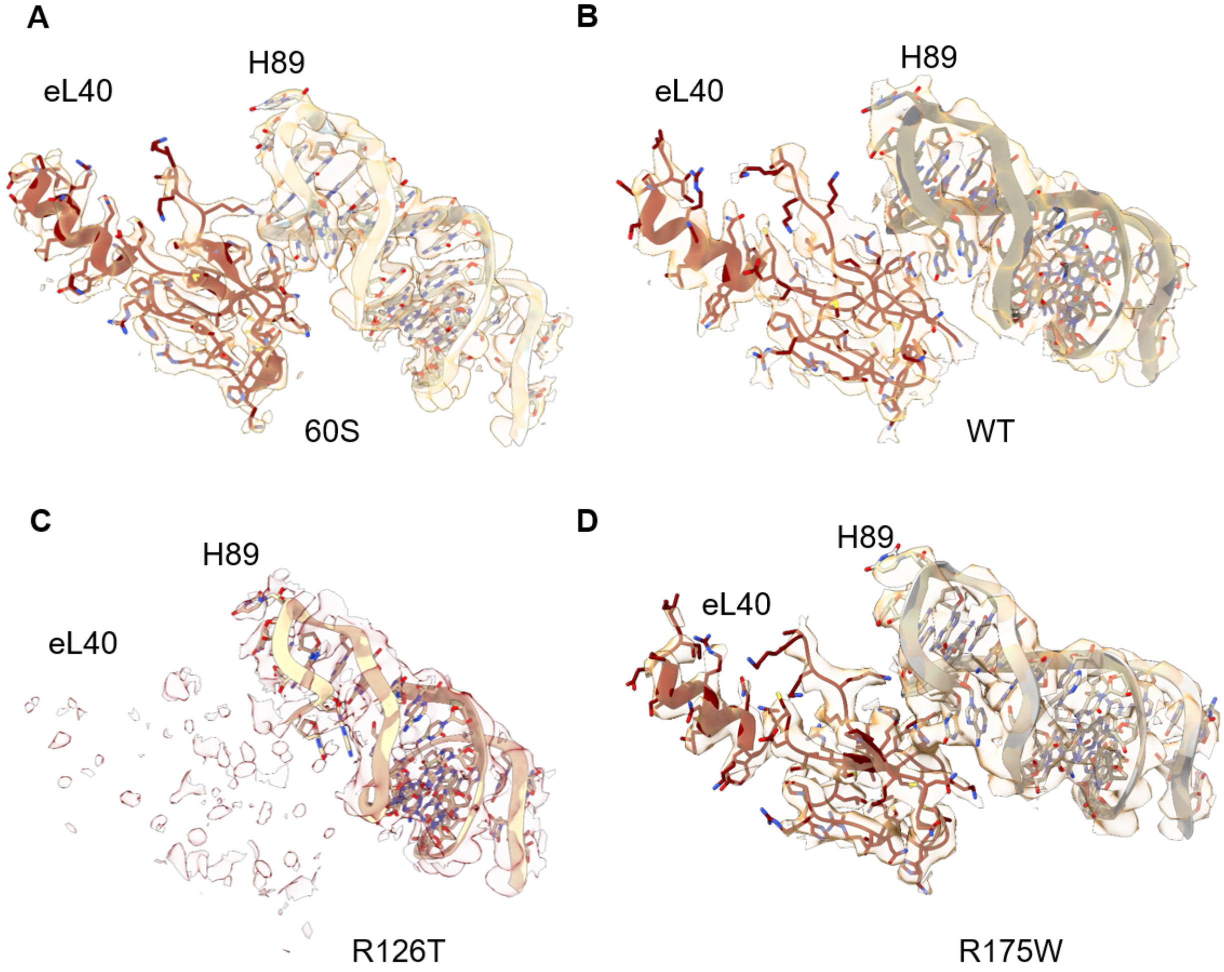


**Supplementary Figure 10. Density for eL40 across the four structures.** Well-ordered density for eL40 is observed in our *(A)* vacant 60S, and 60S structures in complex with *(B)* SBDS-WT, and *(C)* SBDS-R175W. *(C)* In contrast, this density is very sparse in the SBDS-R126T-60S complex. H89 is shown as reference in all four structures and shows excellent density.

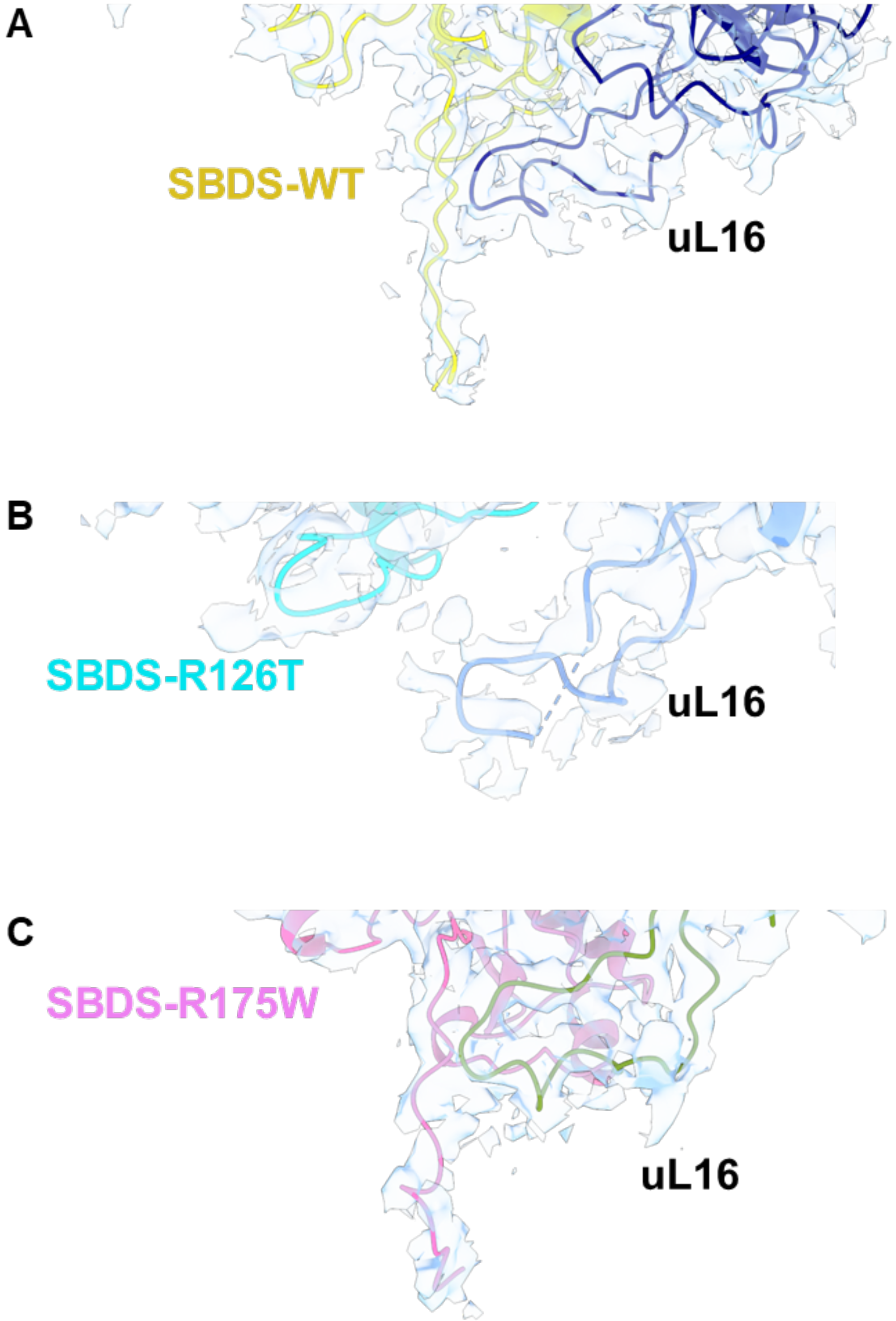


**Supplementary Figure 11. Density in the Cryo-EM data.** Carved density for structures shown in Figure 2E.

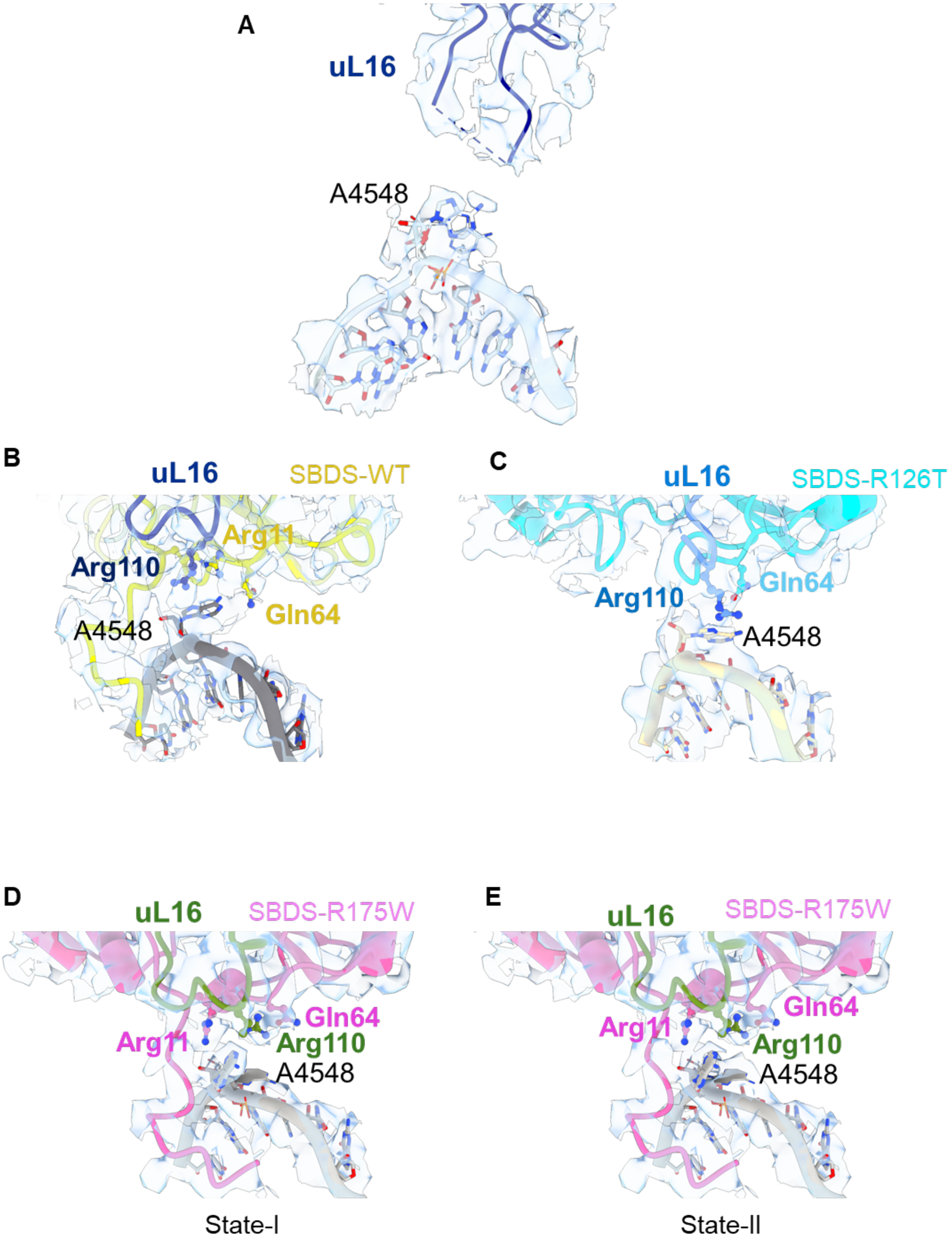


**Supplementary Figure 12. Density in the Cryo-EM data.** Carved density for structures shown in Figure 3. Panels *(A-E)* here correspond to structures in Figure 3, panels A-E.

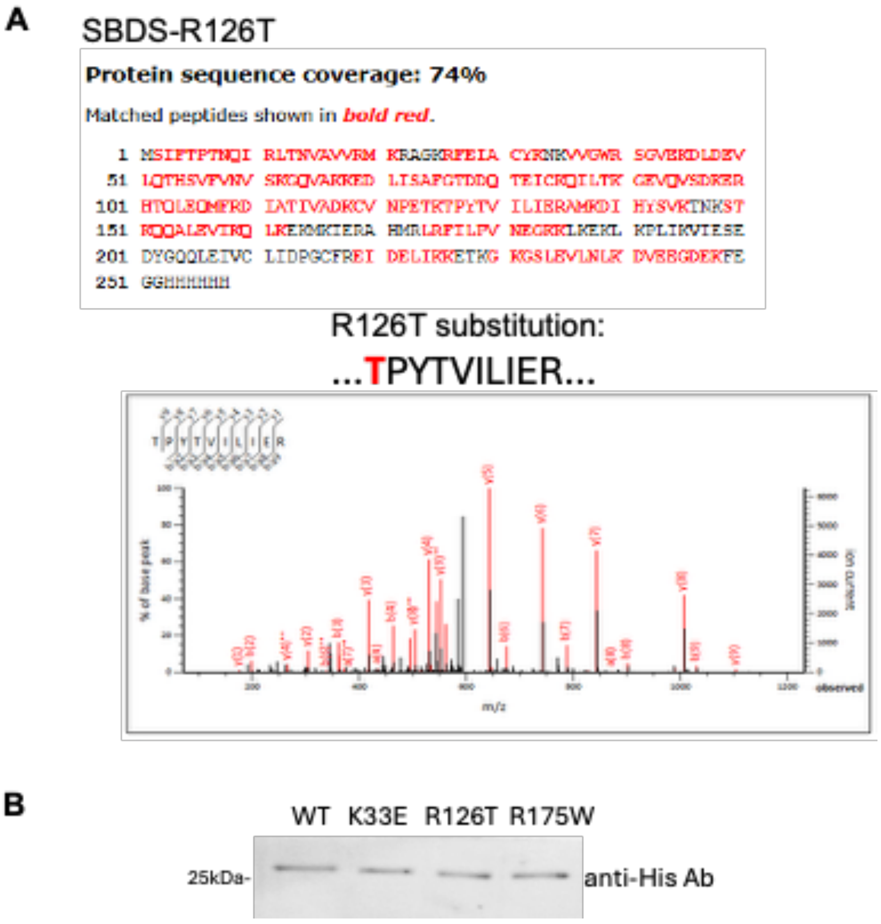


**Supplementary Figure 13. Mass spectrometry analysis of SBDS.** Mass spectrometry of the full-length SBDS-R126T mutant protein confirms presence of an intact N-terminus. Western blot probed with anti-His antibody (SCBT) shows presence of an intact C-terminus in SBDS-WT and all mutant proteins.

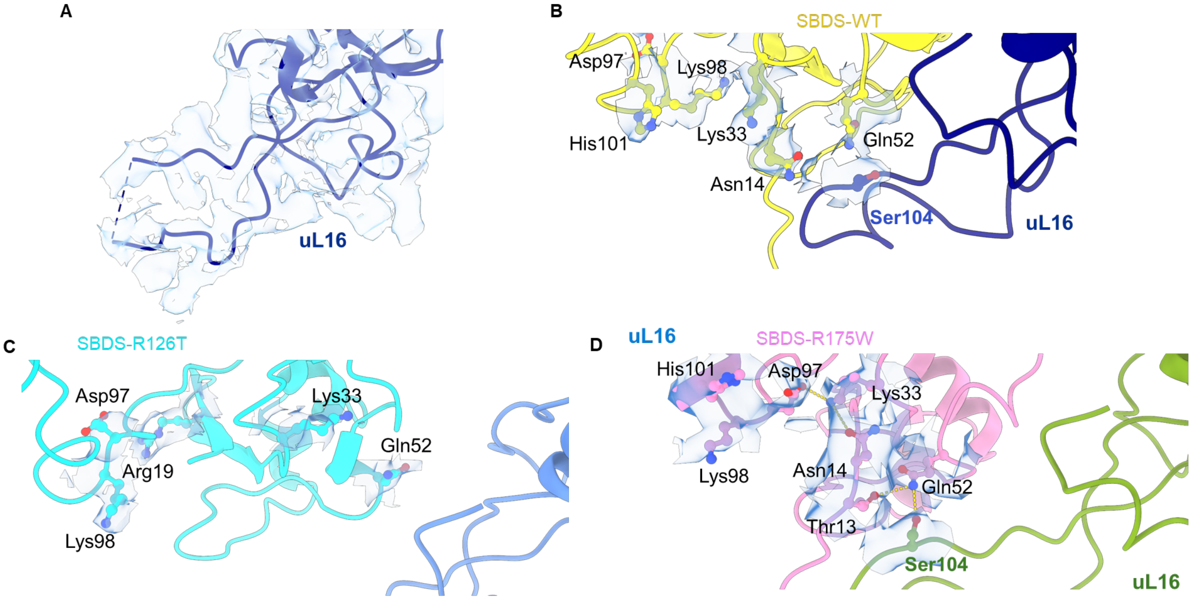


**Supplementary Figure 14. Density in the Cryo-EM data.** Carved density for structures shown in Figure 4. Panels *(A-D)* here correspond to structures in Figure 4, panels A-D.

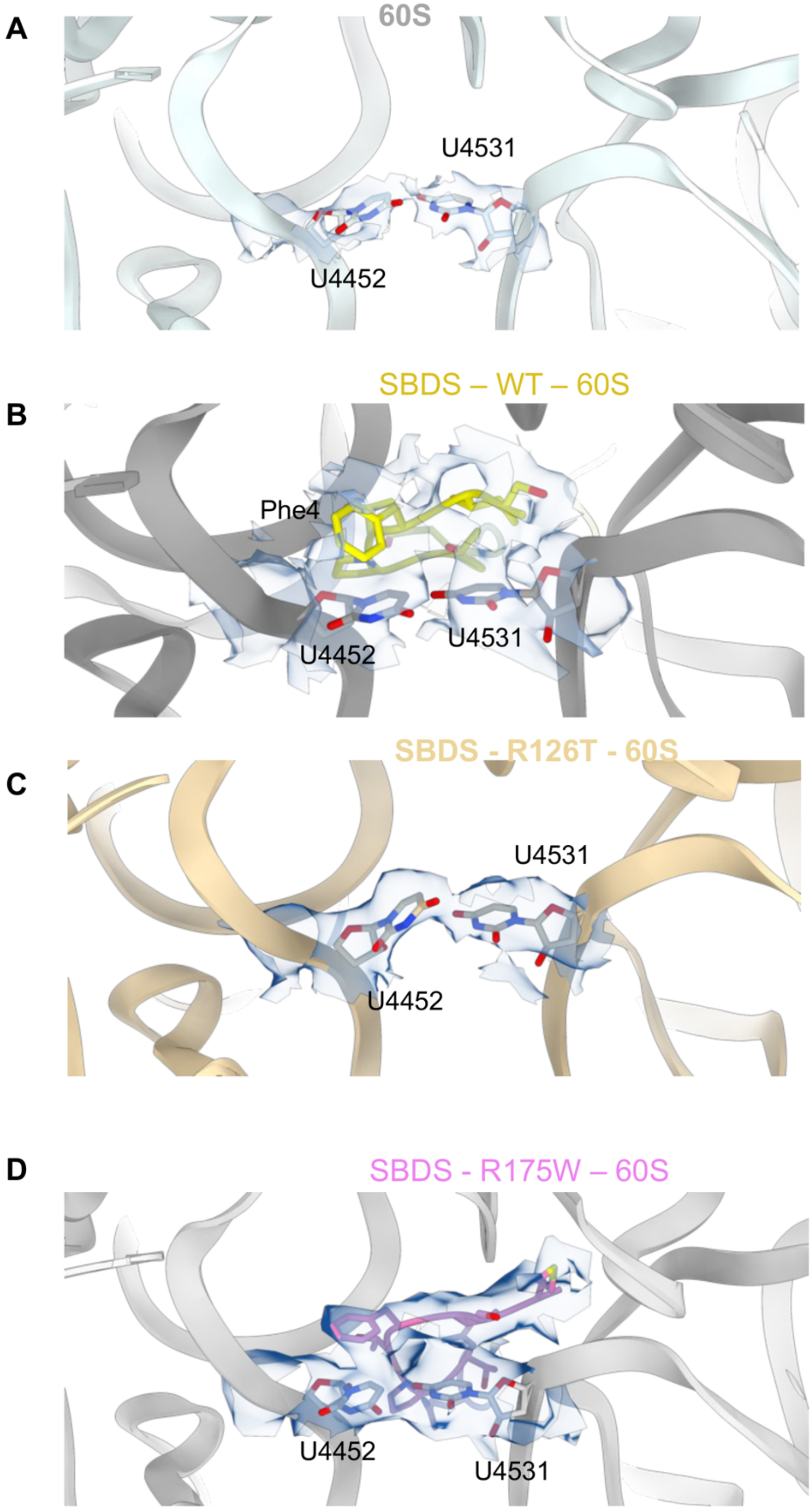


**Supplementary Figure 15. Density in the Cryo-EM data.** Carved density for structures shown in Figure 5. Panels *(A-D)* here correspond to superimposed structures in Figure 5.

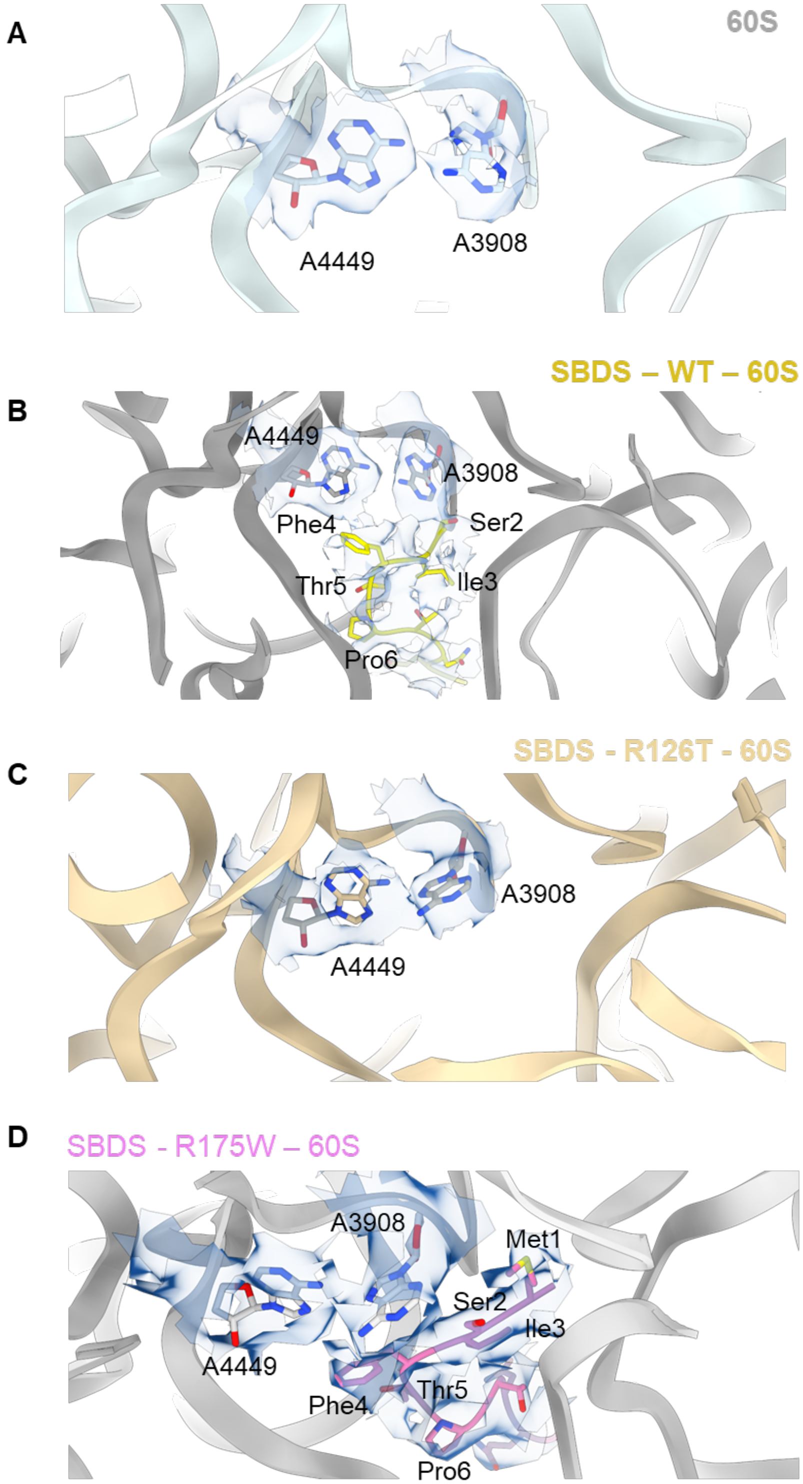


**Supplementary Figure 16. Density in the Cryo-EM data.** Carved density for structures shown in Figure 6. Panels *(A-D)* here correspond to structures in Figure 6, panels A-D.

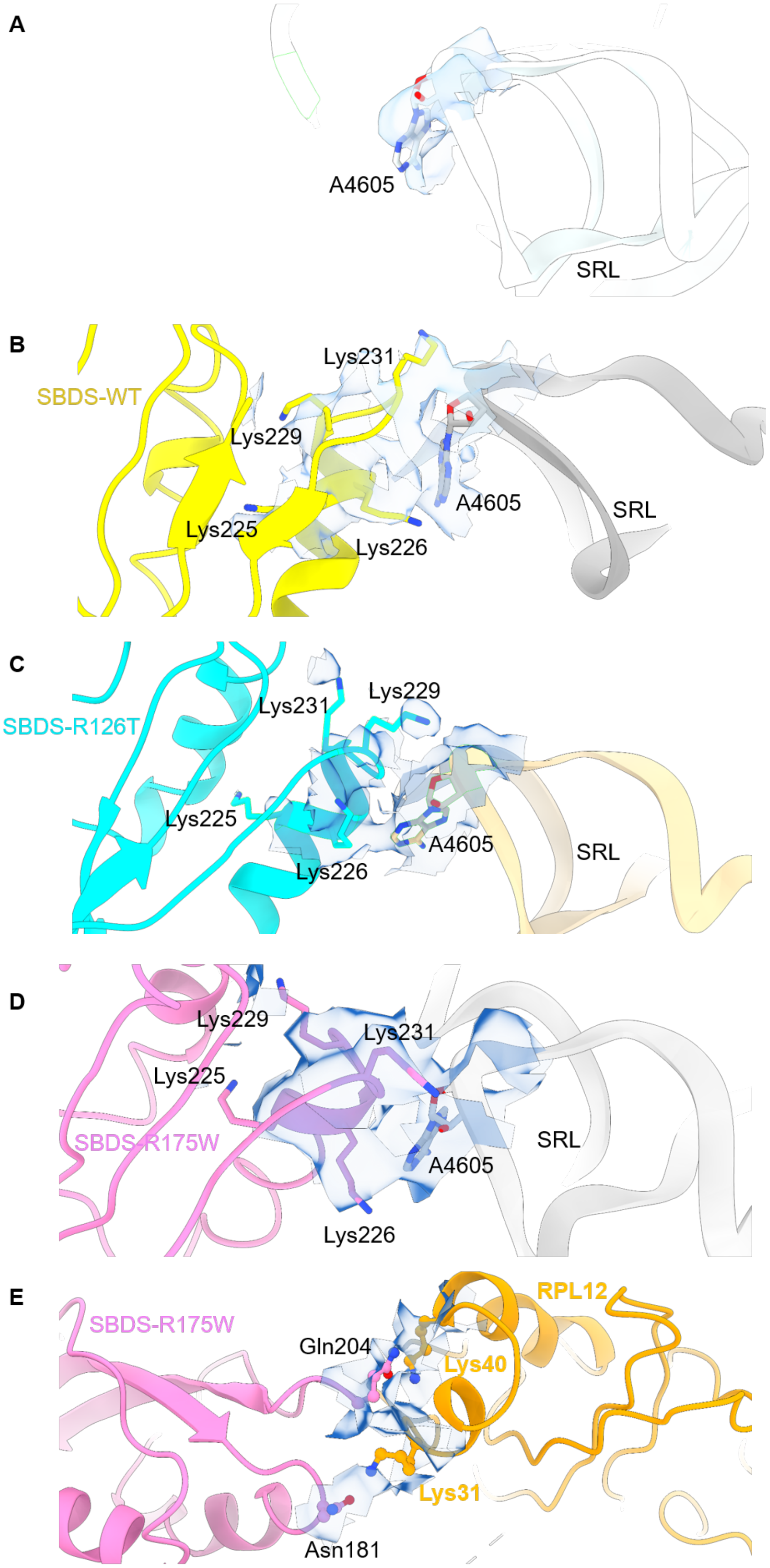
A
A4605
SRL
B
SBDS-WT
Lys231
Lys229
A4605
SRL
Lys225
Lys226
C
SBDS-R126T
Lys231
Lys229
Lys225
A4605
SRL
Lys226
D
Lys229
Lys231
Lys225
SBDS-R175W
SRL
A4605
Lys226
E
SBDS-R175W
RPL12
Gln204
Lys40
Lys31
Asn181

**Supplementary Figure 17. Density in the Cryo-EM data.** Carved density for structures shown in Figure 7. Panels *(A-E)* here correspond to structures in Figure 7, panels A-E.

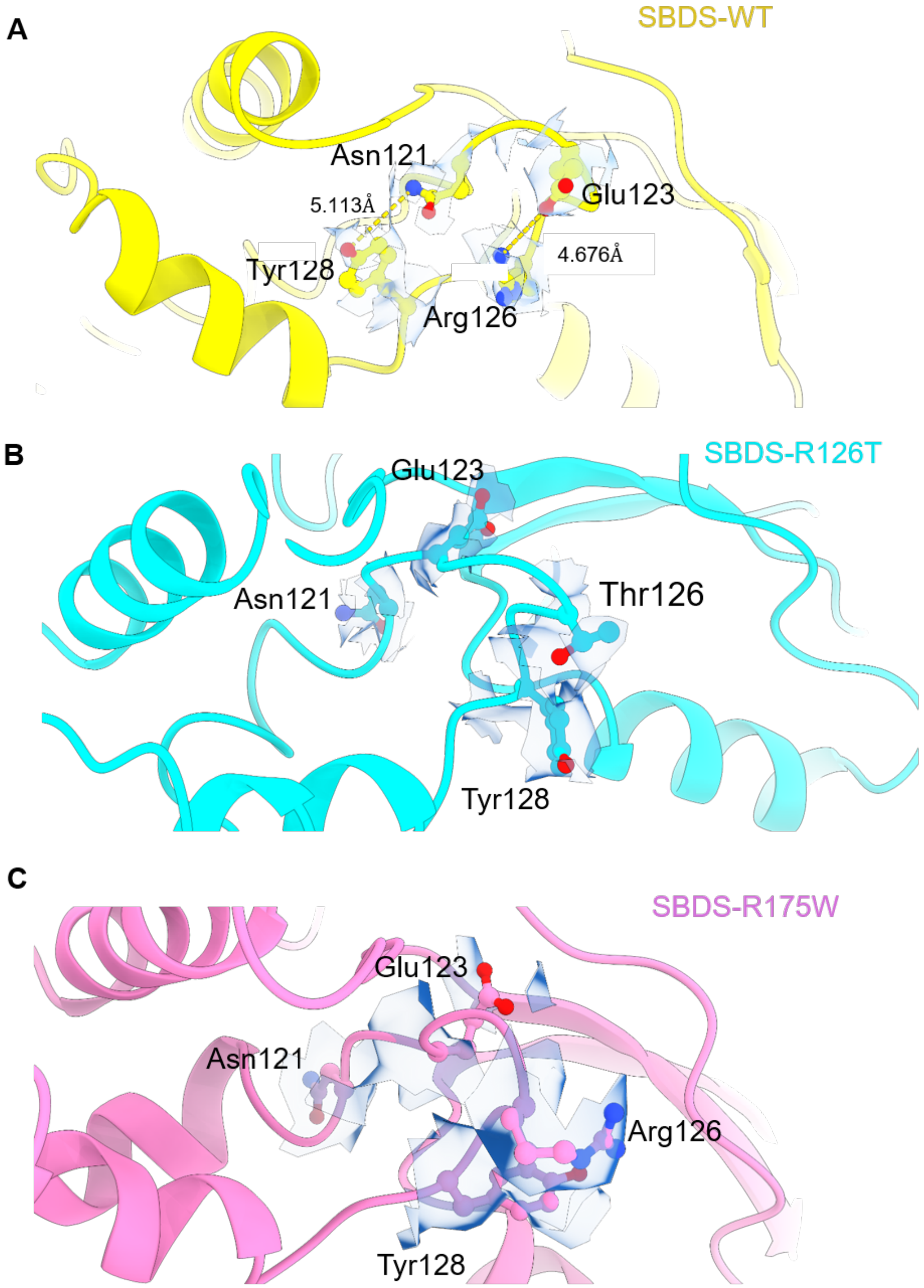


**Supplementary Figure 18. Density in the Cryo-EM data.** Carved density for structures shown in Figure 8. Panels *(A-C)* here correspond to structures in Figure 8, panels A-C.

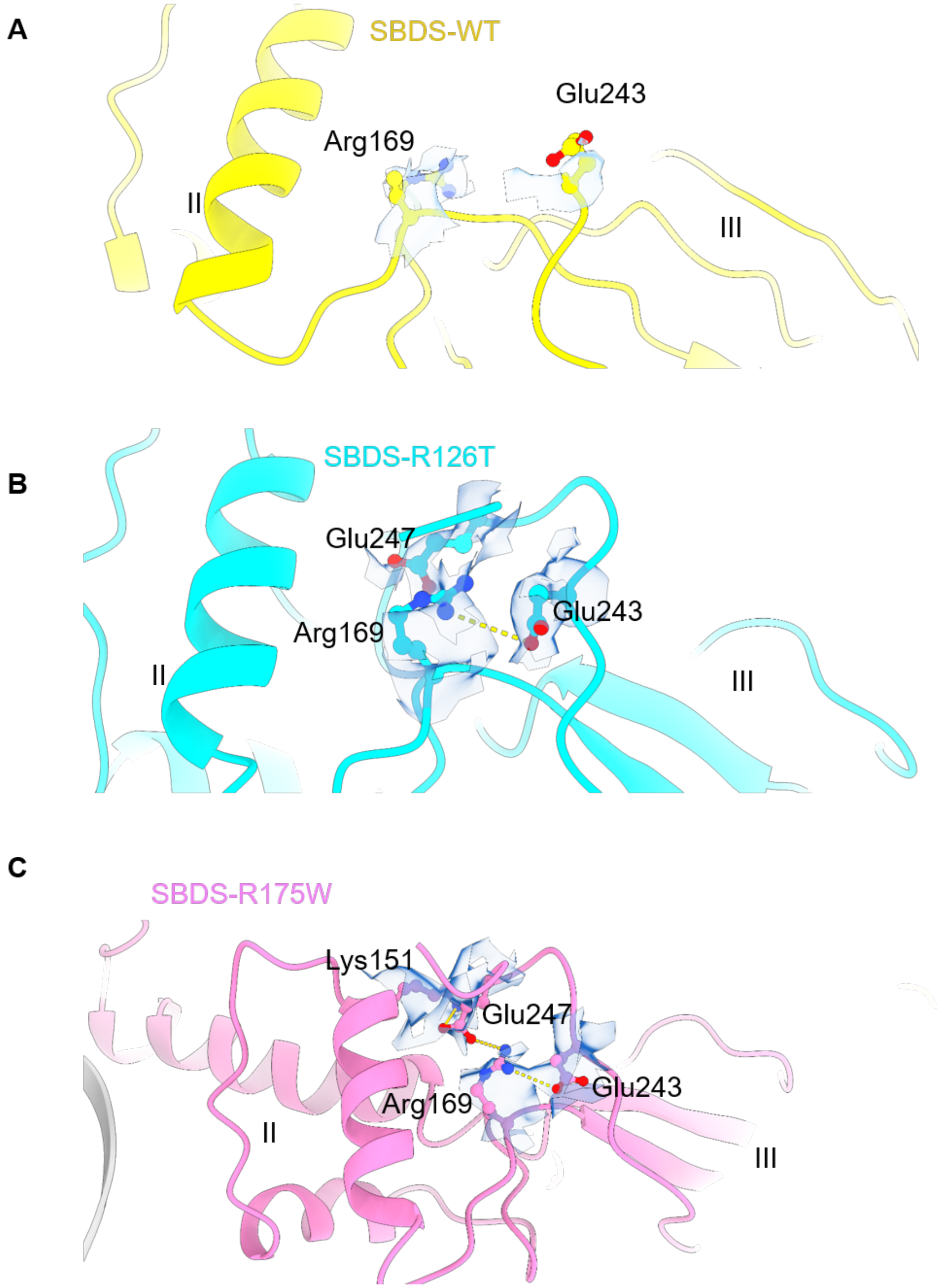


**Supplementary Figure 19. Density in the Cryo-EM data.** Carved density for structures shown in Figure 10. Panels *(A-C)* here correspond to structures in Figure 10, panels A-C.

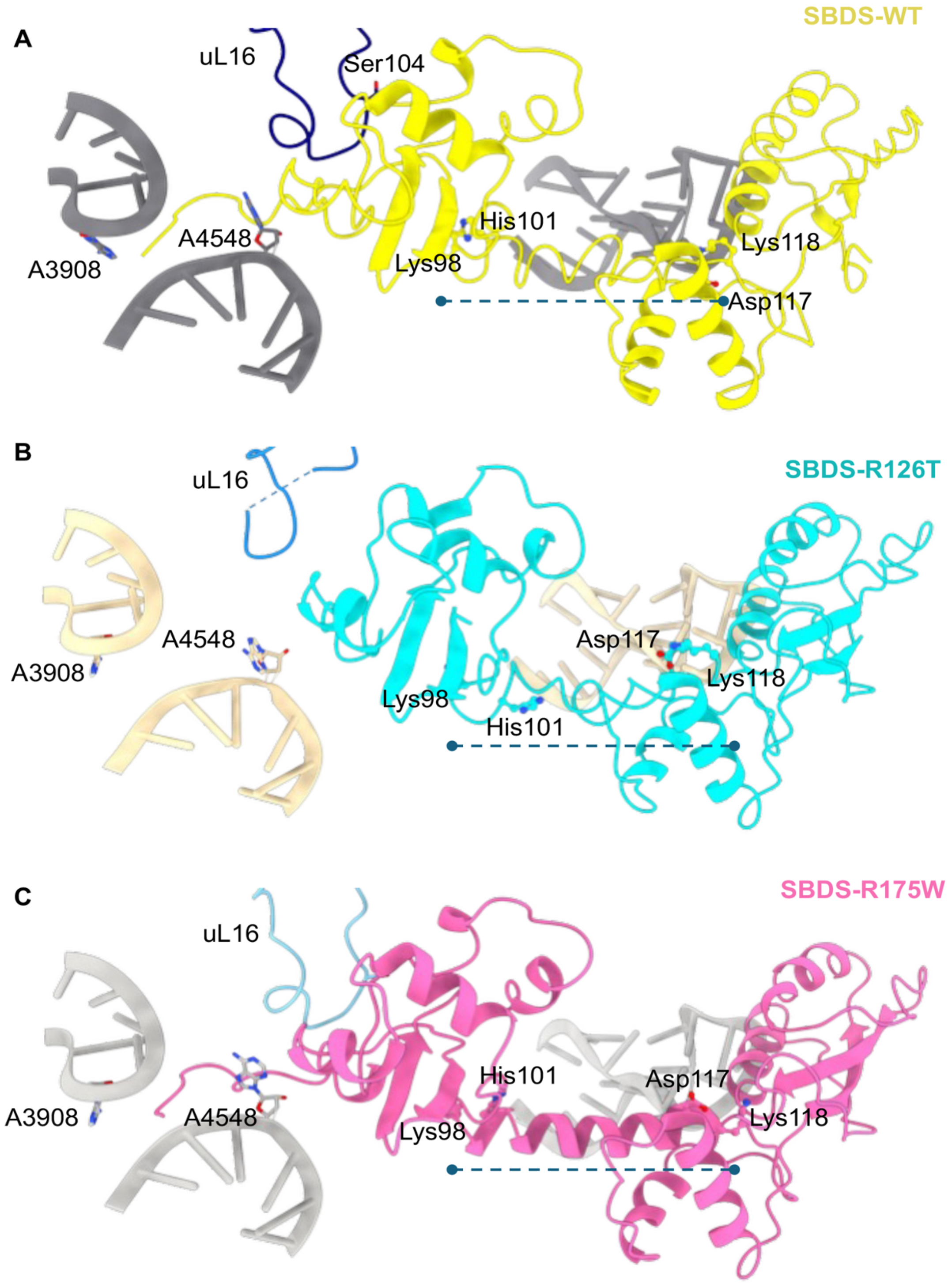


**Supplementary Figure 20. Comparison of changes in the central helix of domain II.** The central helix of domain II connecting domains I and II are denoted by the dashed line in *(A)* SBDS-WT, *(B)* SBDS-R126T, and *(C)* SDBS-R175W. Residues Lys98 and His101 cap one side of the central helix whereas Asp117 and Lys118 cap the other side. These residues undergo large scale rotational movements in the three structures and are driven by allosteric changes introduced by the two respective mutations.

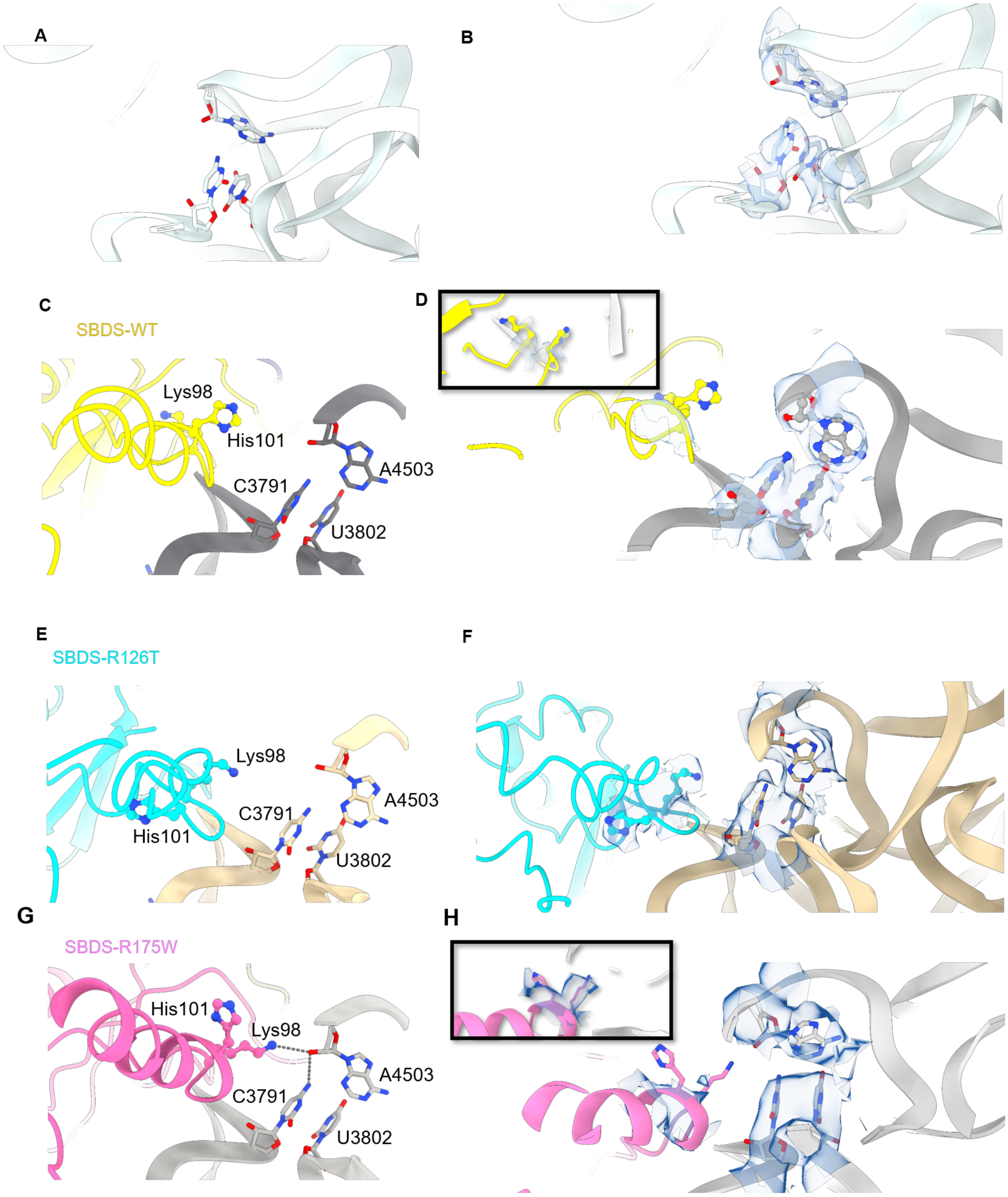


**Supplementary Figure 21. Movement of the His-101 gating residues.** Positions of nucleotides C3791, U3802, and A4503 are shown in *(A,B)* Vacant 60S. Positions of His101 and Lys98 and nucleotides C3791, U3802, and A4503 are shown in *(C,D)* SBDS-WT-60S, *(E,F)* SBDS-R126T-60S, and *(G,H)* SBDS-R175W-60S complexes. While no interactions are observed in the SBDS-WT and SBDS-R126T bound 60S complexes, Lys98 forms a network of interactions with both

C3791 and A4503 in the SBDS-R175W-60S complex. The densities in the Cryo-EM data are shown in the panels on the right.
